# Phosphorylation of a conserved intrinsically disordered region is necessary for activation of a bacterial Hanks-type Ser/Thr kinase signaling pathway

**DOI:** 10.1101/2025.11.01.686061

**Authors:** Noam Grunfeld, Erel Levine, Elizabeth A. Libby

## Abstract

Intrinsically disordered regions (IDRs) are found throughout all domains of life, yet their contribution to bacterial signaling has remained unclear. Here we show that phosphorylation of a conserved juxtamembrane IDR is necessary for activation of the bacterial Hanks-type Ser/Thr kinase-phosphatase PrkC/PrpC signaling pathway in *Bacillus subtilis*. Phosphoablative mutation of a conserved IDR phosphosite has strong effect on kinase-activity-dependent phenotypes, including intrinsic β-lactam resistance and stationary phase survival. Using a synthetic quantitative reporter for kinase activity, mutational analysis, and mathematical modeling, we show that phosphorylation of the IDR promotes trans autoactivation of the kinase, and that this modification is essential for amplifying kinase activity in response to a signal. Phylogenetic analysis demonstrates that this IDR and associated putative phosphosites proximal to the kinase domain are highly conserved across prokaryotic species that diverged at the last universal common ancestor. Together these findings suggest that kinase-domain proximal IDR phosphorylation has a critical role in bacterial Ser/Thr signaling, with direct implications for understanding antibiotic resistance mechanisms and developing kinase-targeting antimicrobials.

**Author Summary:** Bacteria use protein kinases to sense and respond to their environment. Despite decades of research, the activation process of an evolutionarily ancient group of bacterial Ser/Thr kinases that serve as master regulators of responses to antibiotics remains poorly understood. In this work we show that phosphorylation of an unstructured protein region, an intrinsically disordered region (IDR), is required for activating the prototypical Ser/Thr bacterial kinase in *Bacillus subtilis*. Without this phosphorylation, bacteria are more sensitive to β-lactam antibiotics and show severe survival defects in stationary phase. Using genetic analysis and mathematical modeling, we show that IDR phosphorylation allows kinases to activate each other, generating signal amplification. This kinase IDR is conserved across bacteria and archaea that diverged billions of years ago, suggesting it arose early in evolution. Because closely related kinases in clinically important pathogens share these features, our findings suggest this IDR as a target for developing antimicrobials that could disrupt bacterial responses to antibiotic treatment.

## Introduction

Once thought to be restricted to eukaryotes, Hanks-type Ser/Thr kinases^1^ and partner phosphatases are now known to be prevalent in prokaryotes. These systems were first identified in the 1990s based on homology of their catalytic domains to eukaryotic kinases^2^, leading them to be termed eukaryotic-like kinases, or eSTKs. Phylogenetic analysis of their catalytic domains suggests that these systems are evolutionarily ancient, potentially arising at the last universal common ancestor of life on earth (LUCA)^3,4^. Since their initial identification, Hanks-type Ser/Thr kinases have been found to be widespread throughout bacteria and archaea, where they regulate critical cellular processes such as central metabolism, translation, antibiotic sensitivity, virulence, and pathogenesis^5–7^. Among them, the PASTA receptor kinases^8,9^ represent one of the best studied subgroups; they are prevalent throughout Gram positive bacteria, including clinically important pathogens, and can be essential proteins. Their receptors have homology to penicillin-binding domains, suggesting these kinases are involved in sensing cell wall processing^10^. Kinases in this family are master regulators of antibiotic sensitivity and pathogenesis^11^, and consequently, inhibiting their activity has emerged as a potential antimicrobial strategy^12–14^. Despite their importance and clinical potential, we currently lack a mechanistic understanding of bacterial-specific mechanisms for Hanks-type Ser/Thr kinase activation that would facilitate therapeutic development^15^.

To date, most studies on the activation of bacterial Hanks-type Ser/Thr kinases have focused on receptor-ligand interactions. A series of studies in a variety of organisms have shown that for the PASTA kinases, specific residues in the extracellular receptor contact cell wall precursors^10,16–18^, such as muropeptides or lipid II, which function as activating ligands, presumably by inducing kinase dimerization^19,20^. Kinase activation requires phosphorylation, which can occur in trans, as kinases with mutations in the conserved catalytic site (Lys 40 in *B. subtilis* PrkC) can be phosphorylated by active kinases in vitro^21–23^. PrkC is required for *B. subtilis* resistance to cefotaxime^24^, a β-lactam that interferes with cell wall synthesis and likely increases levels of the activating ligand. However, it remains unclear how the signal is transduced from the extracellular receptor to the intracellular catalytic domain to elicit robust downstream signaling.

The PASTA kinases have a conserved intrinsically disordered region (IDR) between the membrane and the catalytic domain, termed the juxtamembrane domain, of generally unknown function^25^. Prior studies identified autophosphorylation sites within these domains but could not connect them to a specific kinase activation mechanism^21,26–28^. For example, phosphoproteomic studies of *B. subtilis* consistently detect the IDR phosphosite at Thr 290 for the prototypical PASTA kinase PrkC^29,30^, demonstrating that kinase IDR phosphorylation occurs in vivo, yet its relevance to physiology or signaling has remained unclear^31,32^.

Here we show that the conserved intrinsically disordered juxtamembrane domain is necessary for activation of the PrkC signaling pathway in *B. subtilis.* We first show that the kinase domain-proximal phosphosite in the IDR, Thr 290, is highly conserved among prokaryotes. We next show that a phosphoablative mutation at this site results in strong physiological phenotypes including stationary phase lysis, antibiotic sensitivity, and downstream changes in gene expression. Using defined titrations of kinase IDR variants and a mathematical model, we demonstrate that this phosphosite is required for full trans-activation of the kinase, activation of the signaling pathway, and ligand-mediated upregulation. Together with the conservation of the IDR and the associated phosphosite in clinically important pathogens, this identifies this region as an attractive drug target.

## Results

### Conserved intrinsically disordered regions with a phosphorylation site-centered motif in Ser/Thr kinases

One understudied common feature of prokaryotic Ser/Thr kinases is the presence of an intrinsically disordered region (IDR) between the kinase domain and the transmembrane domain or a cytoplasmic receptor (bacteria, Fig. 1A; archaea, Fig. S1A). The prototypical *B. subtilis* kinase PrkC has a ∼61 residue intrinsically disordered juxtamembrane (JM) domain between the kinase domain and the transmembrane domain (Fig. 1B). Since IDRs are less common in prokaryotes than eukaryotes^31,33,34^, we hypothesized that this JM domain may have important conserved features. To test this hypothesis, we aligned kinases that were previously used to study phylogenetic conservation of the catalytic domains across bacterial and archaeal species^3^ and extended the analysis to the IDRs. Using AlphaFold structure prediction, we assessed disorder by identifying regions with low to very low per-residue confidence scores (pLDDT < 70, Fig. S1B), which are commonly associated with intrinsic disorder^35,36^. Based on this criterion, we found that all kinases in this group contain an IDR directly proximal to the kinase domain. Among the bacterial kinases with an identifiable transmembrane domain (18/20), the IDR resides in a juxtamembrane domain, ranging in length between 39 and 81 residues. Moreover, sequence alignment revealed a highly conserved region around a Thr/Ser located ∼20 residues from the kinase domain, aligning with the phosphosite Thr 290 in *B. subtilis* (Fig. 1A, top row). This region is strikingly conserved even among evolutionarily distant species (∼3000 MY^37^), suggestive of a functional role.

**Figure 1:**
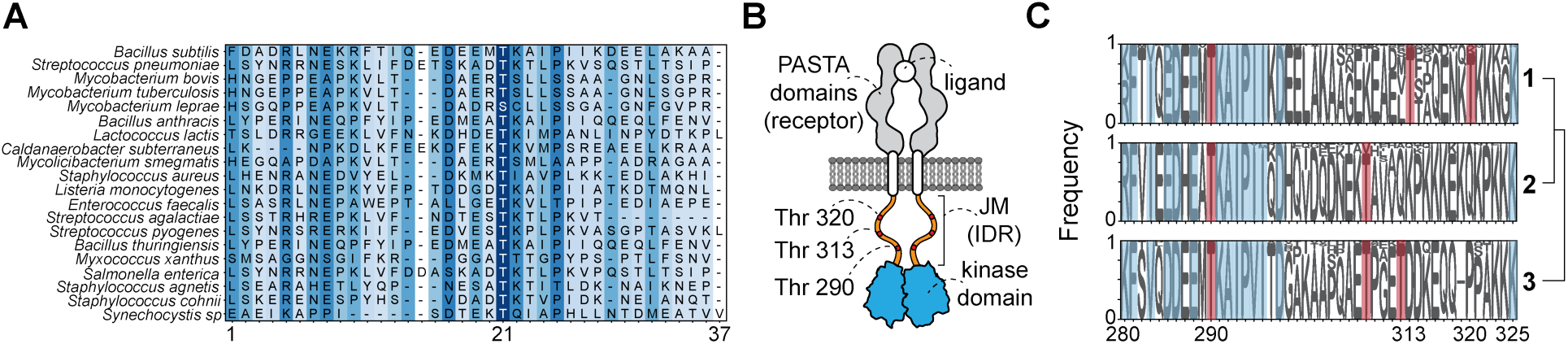
Conservation of the Hanks-type Ser/Thr kinase juxtamembrane IDRs. **A) Multiple sequence alignment.** Kinases from evolutionarily distant species exhibit a kinase-domain proximal IDR with a highly conserved Thr residue. Alignment is shown relative to *B. subtilis* PrkC (top row), with shading indicating the most highly conserved positions (dark blue) and variable residues (light blue to white). **B) Schematic of *B. subtilis* PrkC domain architecture.** PrkC contains three extracellular PASTA domains (grey), an intrinsically disordered juxtamembrane region (orange), and a cytoplasmic kinase domain (cyan). The juxtamembrane IDR has three established phosphothreonine sites (red). In the presence of an activating ligand (white), PrkC is hypothesized to exhibit ligand-binding-mediated dimerization. **C) Sequence conservation of IDRs from PrkC homologs across *Bacilli*.** Kinase homologs from 500 *Bacillus* genomes were aligned and grouped into three hierarchical clusters, indexed relative to *B. subtilis* PrkC, cluster 1. Known (Thr 290, 313, and 320) and potential (Thr 308 and 312) phosphothreonines are highlighted in red. The other residues most highly conserved across all clusters are highlighted in blue. Thr 290 and its surrounding residues show the strongest conservation.

In vitro studies previously identified three phosphothreonine sites in the PrkC IDR, the highly conserved site Thr 290, and two additional sites: Thr 313 and Thr 320^21^ (Fig. 1B). To examine the conservation of these phosphosites and their surroundings, we aligned IDR sequences from 500 homologs of *B. subtilis* PrkC (Materials and Methods). All 119 unique sequences show significant conservation surrounding the Thr 290 phosphosite, particularly through the +6 position relative to this site (Fig. 1C). This suggests conservation of the phosphosite, since recognition and phosphorylation of substrates by Ser/Thr kinases is governed by the surrounding residues^38,39^. The remainder of the IDR shows relatively weaker conservation.

Based on IDR sequence, 464 of the 500 species can be placed into 3 phylogenetic groups. Within each group, the entire IDR shows little sequence variation and has potentially one or two additional phosphothreonines by homology to the known sites. However, the positioning of these sites and the relative conservation of their surrounding residues vary. All IDR sequences have several lysine residues adjacent to the transmembrane domain, interspersed with other positively charged hydrophilic residues, suggesting a potential function in preventing membrane association^40^. Together, these observations reveal that an intrinsically disordered JM domain with at least one conserved phosphosite, Thr 290 in *B. subtilis*, is a widespread feature of bacterial Hanks-type Ser/Thr kinases.

### The highly conserved IDR phosphosite is necessary for PrkC-dependent phenotypes in vivo

Given the strong conservation of the IDR and its phosphosites, we sought to characterize their functional importance for kinase activity in vivo. We assessed the importance of these sites to cellular physiology using two kinase activity-dependent phenotypes: stationary phase lysis^41^ and antibiotic sensitivity^24^. The importance of the conserved threonine phosphosites in the IDR was tested using allelic replacements of *prkC* at the native chromosomal locus with known phosphoablative Thr-to-Ala mutations^21^. Allelic replacements were verified by both direct sequencing of the locus and whole genome sequencing (Materials and Methods).

First, we examined the effect of phosphoablative IDR mutations on PrkC activity-dependent colony lysis on agar plates (Fig. 2A). We observed faster onset of lysis in a *prkC^T290A^* background than in *prkC^WT^*. This effect was comparable to the lysis observed in both the *prkC^K40A^*, a catalytic site mutation^17,42^, and the Δ*prkC* deletion backgrounds, suggesting that the phosphoablative mutation T290A has a strong effect on *B. subtilis* physiology that is comparable to complete inactivation of the kinase. In contrast, for the two other phosphoablative IDR mutants, *prkC^T313A^* and *prkC^T320A^*, the phenotype appeared similar to *prkC^WT^*, suggesting that phosphorylation of those IDR residues does not play a significant role in the context of lysis.

**Figure 2:**
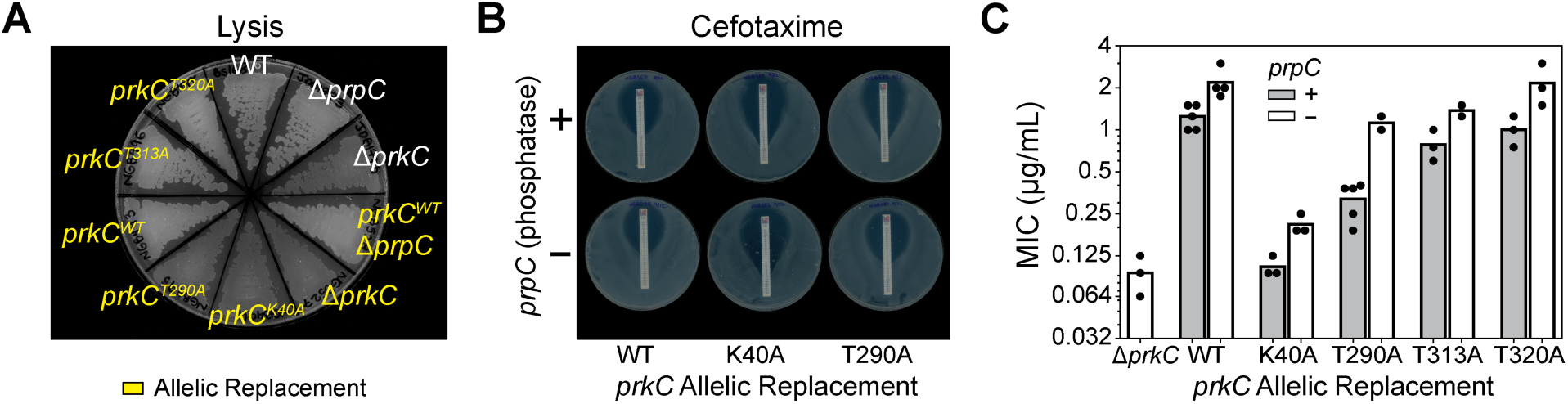
PrkC IDR phosphosite-dependent phenotypes: lysis and β-lactam sensitivity. **A) IDR phosphosite-dependent lysis on agar plates**. Growth and lysis of kinase (*prkC*) and phosphatase (*prpC*) mutants. White: wild-type *B. subtilis* 168 and previously published mutants. Yellow: allelic replacement strains, harboring a catalytic site mutation, *prkC^K40A^*, phosphoablative mutations of the conserved Thr 290 phosphosite, *prkC^T290A^*, or two additional sites, *prkC^T313A^* and *prkC^T320A^*. Lysis appears darker grey than robust growth. **B) IDR phosphosite-dependent sensitivity to the β-lactam cefotaxime.** Representative antibiotic test-strip sensitivity assays comparing *prkC^WT^, prkC^K40A^,* and *prkC^T290A^* allelic replacements in otherwise wild-type (+) and phosphatase knockout Δ*prpC* (-) backgrounds. Intersection of the bacterial lawn with the test strip indicates the MIC. **C) Quantification of sensitivity assays shown in B.** Bars represent mean measured MICs across biological repeats (dots, n = 2-5). *prkC^K40A^* and *prkC^T290A^* reduce MIC 12-fold (±1.6) and 4-fold (±0.6), respectively, compared to *prkC^WT^*. Deletion of the phosphatase (Δ*prpC*) restores the MIC of *prkC^T290A^* to near wild-type levels.

As in many homologous systems, *B. subtilis* PrkC regulates sensitivity to the β-lactam cefotaxime by approximately 10-fold^24,43–46^. Since the lysis phenotype in a *prkC^T290A^* background is consistent with a loss of kinase activity, we hypothesized that this mutation would increase cefotaxime sensitivity and be measurable in a standard antibiotic test strip assay (Fig. 2B). In this assay, a test strip containing a gradient of cefotaxime is placed on an agar plate inoculated with a confluent lawn of bacteria. The intersection of bacterial growth with the strip indicates the minimum inhibitory concentration (MIC) of the antibiotic. We found that a *prkC^WT^*allelic replacement strain had a similar MIC to the *B. subtilis* 168 WT parent strain (∼1.25 µg/ml), consistent with published results^24^ (Fig. 2C). In contrast, a catalytic mutant *prkC^K40A^* allelic replacement had a significant increase in sensitivity (MIC ∼0.1 µg/ml), similar to a kinase deletion (Δ*prkC*). The *prkC^T290A^* allelic replacement resulted in a MIC of ∼0.3 µg/ml, approximately 4-fold more sensitive than *prkC^WT^*. However, in a phosphatase knock-out background (Δ*prpC*), the T290A mutation did not have a strong effect, while the K40A mutation and kinase deletion both maintained a strong increase in sensitivity. This suggests that phosphoablation of Thr 290 reduces the activity of PrkC but does not abolish it. If this is the case, the residual activity of this mutant is enough to support intrinsic resistance to cefotaxime in the absence of the phosphatase, but not if competing with its activity. As in the lysis assay, the cefotaxime sensitivities of the allelic replacement strains *prkC^T313A^* and *prkC^T320A^* were comparable to *prkC^WT^*, and those sites will not be further studied here. Together, these data demonstrate that the most highly conserved phosphosite within the IDR, Thr 290, has a significant yet previously unappreciated role in two important kinase activity-dependent phenotypes.

### IDR phosphorylation affects PrkC-dependent gene expression

Considering the physiological effects observed in a phosphoablative *prkC^T290A^* background, we hypothesized that this mutation would result in quantifiable changes in downstream gene expression. One of the best characterized PrkC-dependent effects on transcription is through phosphorylation of WalR, a master regulator of cell wall metabolism^47–49^. PrkC-dependent phosphorylation of WalR at Thr 101 strengthens its regulation of downstream targets^50^, including increasing repression of *pdaC* (*yjeA*)^51^, a cell-wall-associated peptidoglycan deacetylase, and *iseA* (*yoeB*)^52^, a secreted protein involved in cell wall stress response. To quantify the effect of PrkC IDR mutations on *iseA* and *pdaC* expression, we used luminescence-based transcriptional reporters (Fig. 3A, S2A). In a *prkC^T290A^* background, the expression of both genes was significantly upregulated, approximately 4-fold for *iseA* (Fig. 3B) and 2-fold for *pdaC* (Fig. S2B) compared to a *prkC^WT^* background. This upregulation is similar to that observed in a *prkC^K40A^* or a kinase deletion background, Δ*prkC*, demonstrating that this IDR phosphosite is likely required for full kinase activity and downstream signaling.

**Figure 3:**
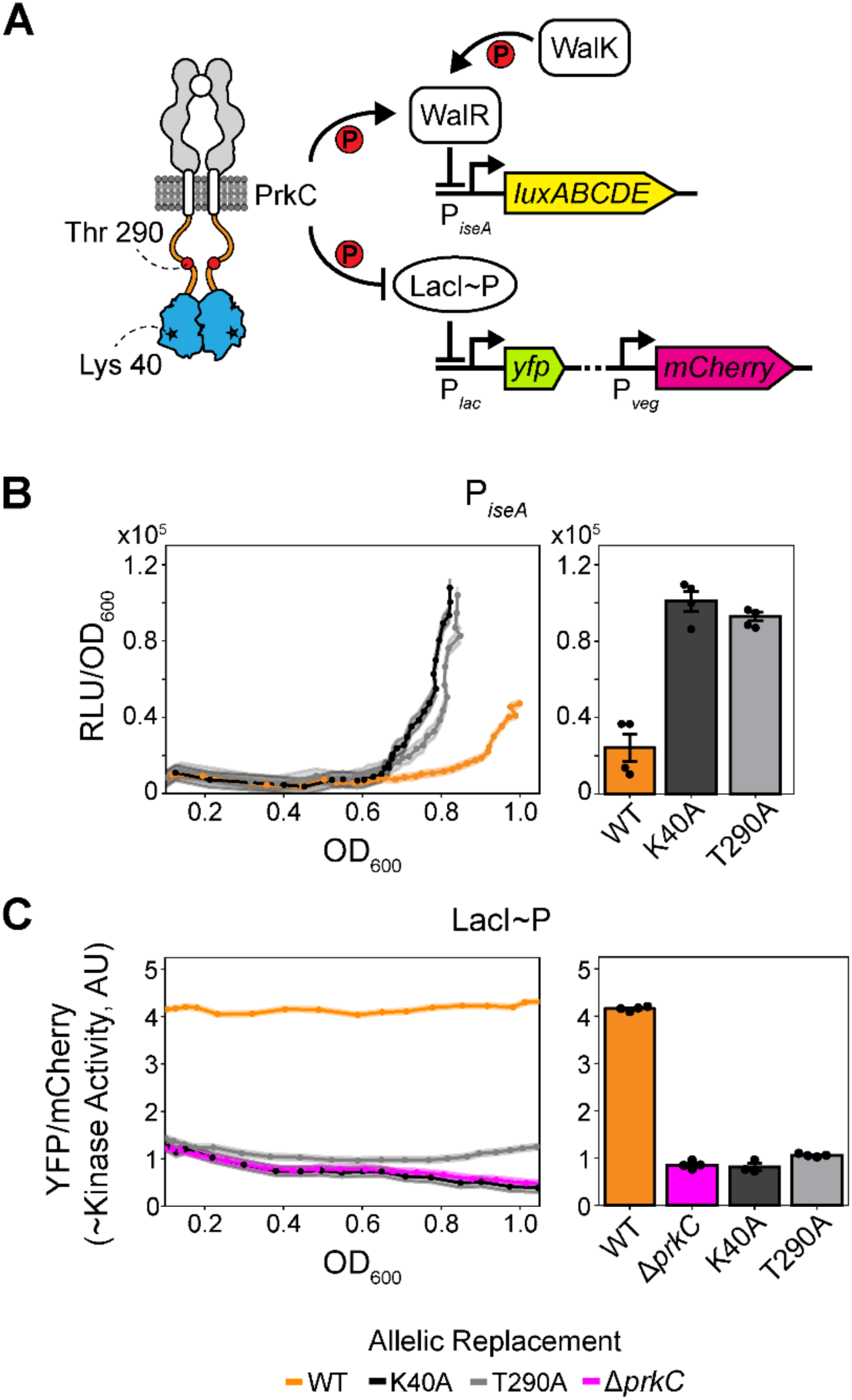
Transcriptional reporters of PrkC activity show Thr 290 phosphosite-dependent changes. **A) Schematic of native and synthetic transcriptional reporters for PrkC activity. Top:** Native reporter system. PrkC and WalK phosphorylate WalR, increasing its repression of WalR-regulon genes such as *iseA* in stationary phase. **Bottom:** Synthetic reporter system to isolate PrkC activity. PrkC phosphorylates LacI∼P, a synthetic transcription factor, relieving repression of *yfp*. Constitutively expressed *mcherry* is used for normalization. YFP/mCherry serves as a proxy for total PrkC kinase activity. **B) A WalR-regulon transcriptional reporter shows Thr 290-dependent regulation. Left:** Normalized luminescence (RLU/OD_600_) as a function of growth (OD_600_) for the P*_iseA_-lux* reporter in *prkC^WT^* (orange), *prkC^T290A^* (grey), and *prkC^K40A^*(black) allelic replacement strains in a *prpC^+^* background. Dots show means of 3 replicates, shading indicates standard deviation. ***Right:*** Bar plot shows means at early stationary phase. Error bars indicate the SEM from ≥4 independent experiments (dots, n = 4-5), including the representative experiment shown on the left. Both *prkC^T290A^* and *prkC^K40A^* mutants show a 4-fold (±1) increase in signal. **C) A synthetic reporter system for PrkC activity shows Thr 290 phosphosite-dependent regulation. Left:** Kinase reporter activity (YFP/mCherry) as a function of growth (OD_600_) in *prkC^WT^* (orange), *prkC^T290A^* (grey), *prkC^K40A^* (black), and Δ*prkC* (magenta) allelic replacement strains in a *prpC^-^* background. Dots represent means of 3 replicates and shading indicates standard deviations. **Right**: Bar plot of mean normalized fluorescence in log phase. Error bars represent the SEM from ≥3 independent experiments (dots, n = 3-4), including the representative experiment shown on the left. The *prkC^T290A^* and *prkC^K40A^* mutants result in 3.9-fold (± 0.1) and 5.1-fold (± 0.5) reductions in signal, respectively, comparable to Δ*prkC*.

### A system for quantitative analysis of IDR-dependent PrkC activation

Measuring PrkC activity using WalR-dependent changes in gene expression is complicated by the known presence of additional regulators of WalR, including its essential partner histidine kinase WalK (Fig. 3A). To overcome this complexity, we used a synthetic reporter for kinase activity that is comparatively insulated from the native transcriptional networks (Fig. 3A). This system uses a phosphorylation-sensitive transcription factor, LacI∼P, to regulate a fluorescent transcriptional reporter^24^. LacI∼P is composed of a phosphopeptide binding domain (FHA2) translationally fused to the *lac* repressor LacI and to a short phosphosubstrate peptide which is a known target of the kinase. PrkC-dependent phosphorylation of this transcription factor results in downstream expression of a fluorescent reporter (*yfp*). For normalization, a second fluorescent reporter (*mCherry*) is expressed constitutively. This system has been shown to be less sensitive to growth phase and media than the WalR regulon^53^, suggesting it is a more direct measure of PrkC activity. To isolate the effect of IDR mutations on the kinase, we used this reporter to measure activity in the absence of the phosphatase (a Δ*prpC* background). We observed a comparable loss in activity in *prkC^T290A^*, *prkC^K40A^*, and Δ*prkC* backgrounds of ∼4.5-fold compared with *prkC^WT^* (Fig. 3C).

Our phenotypic (Fig. 2) and transcriptional (Fig. 3) data indicate that phosphorylation of the juxtamembrane IDR at Thr 290 is crucial for kinase activation. Although the function of juxtamembrane domains in prokaryotic Hanks-type Ser/Thr kinases remains unclear, kinase cis-regulatory domains are widespread across evolution^54,55^. Therefore, we sought to characterize the role of IDR phosphorylation in kinase activation, and to determine the mechanism involved.

To this end, we aimed to develop an experimentally supported quantitative model of PrkC activity and its modulation by the Thr 290 phosphosite in the IDR. Quantitative models have been valuable in uncovering activation mechanisms in bacterial signaling systems^56–63^. To do so, they often use measurements of dose-response relationships between the input (stimulus, kinase activity), the intermediate (transcription factor phosphorylation), and the output (reporter gene expression). In the absence of such measurements, we hypothesized that a model for IDR function could be developed based on the quantitative response of the LacI∼P system, a proxy for total kinase activity, to defined titration of kinase variants. We created a system for the heterologous induction of *prkC* IDR variants, expressed from a single copy chromosomal location (*ganA*::P*_xyl_*-*prkC*). To avoid confounding effects of the phosphatase, we first consider experiments in a phosphatase knockout background (Δ*prpC*) and examine the impact of PrpC on kinase activation later in this paper.

We calibrated our system to translate amounts of xylose induction into units of PrkC levels, relative to those observed in the native system (Materials and Methods, Fig. S3). Our mapping assumes that *prkC* expression is not under the feedback control of PrkC activity, an assumption that is supported by our data and discussed further below.

### PrkC autophosphorylation results in positive feedback

To develop a quantitative model for PrkC activation, we measured fluorescence from the LacI∼P reporter at different expression levels of PrkC and several PrkC variants. Titration of different variants revealed distinct activation behaviors (Fig. 4A). While kinase activity responded nearly linearly to increasing expression of PrkC^WT^ (Fig. 4A, orange), titration of PrkC^T290A^ resulted in a non-linear response: activity remained low until expression surpassed a threshold, beyond which it increased almost linearly (Fig. 4A, grey). Titration of the catalytic mutants PrkC^K40A^ and PrkC^K40A^ ^T290A^ resulted in no detectable kinase activity at any tested expression level (Fig. 4A(i), black and red).

**Figure 4:**
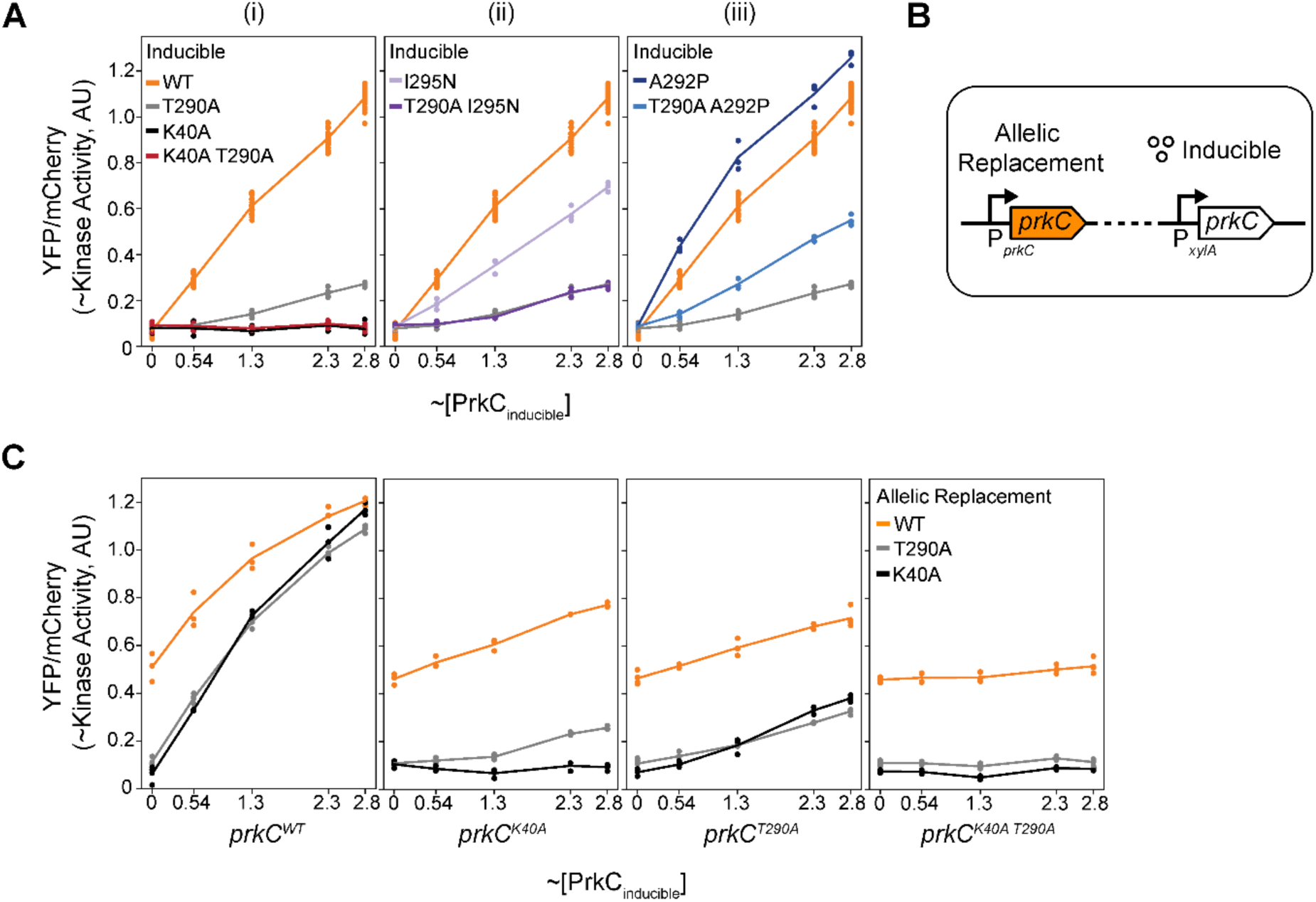
Titration of *prkC* variants shows IDR-dependent regulation of kinase activation. **A) i) Quantification of residual kinase activity of PrkC variants**. Dose response of kinase activity of *prkC^T290A^*, *prkC^WT^*, and catalytic mutants *prkC^K40A^* and *prkC^K40A^ ^T290A^*, titrated at ∼0-2.8x native PrkC levels. The dose-response data were used as the basis for a quantitative model for positive feedback in activation and threshold-like behavior (SI Text). **ii & iii) Residues surrounding the phosphosite affect Thr 290-dependent activation.** **ii) A PrkC^I295N^ variant results in a Thr 290-dependent loss of kinase activity.** Titration of *prkC^I295N^* and *prkC^T290A^ ^I295N^* compared with *prkC^WT^* and *prkC^T290A^* (data repeated from i). **iii) A PrkC^A292P^ variant results in a reduction of Thr 290-dependent dynamic range in kinase activity.** Titration of *prkC^A292P^*and *prkC^T290A^ ^A292P^* compared with *prkC^WT^*and *prkC^T290A^* (data repeated from i). **B) Schematic of kinase variant co-expression experiments to measure the role of Thr 290 in inter-kinase interactions and activation.** In co-expression experiments, one *prkC* variant is under the P*_prkC_* promoter at the native locus; another is under a P*_xylA_* inducible promoter integrated at the *ganA* locus. **C) Measurement of total kinase activity of the co-expression system.** Kinase activity of co-expression strains with allelic replacements at the native locus containing *prkC^WT^*, *prkC^K40A^*, or *prkC^T290A^*, in combination with inducible expression of the kinase variant indicated on each panel. For all panels, each dot represents the mean YFP/mCherry of 3 replicates from biologically independent experiments (WT n = 19-24 in A, n = 3-5 for all else in A and C).

One possible mechanism behind the non-linearity observed in PrkC^T290A^ activity could be positive feedback in kinase autophosphorylation, either in cis or in trans. Consistent with prior modeling of similar systems^56,64^, a simple mathematical model of PrkC activation (SI Text) shows that autophosphorylation alone is sufficient to generate the observed thresholding behavior: kinase activity remains low until a critical total concentration is reached, beyond which the phosphorylated active form accumulates rapidly. Notably, this behavior can arise without requiring changes in kinase dimerization state. This explains why the T290A variant requires higher expression to sustain activation, as the IDR phosphoablative mutation effectively raises the activation threshold for the kinase. As discussed below, this model also suggests that the catalytic mutants appear inactive because their activation threshold exceeds the tested concentrations, not because they cannot activate.

### Conserved residues enhance phosphorylation-dependent changes in kinase activation

The sequence near Thr 290 is highly conserved, particularly in the residues immediately surrounding the phosphosite (Fig. 1A,C), suggesting that both the phosphosite and its local context are important for kinase function. To test this hypothesis, we introduced mutations in two highly conserved positions, Ala 292 and Ile 295, which may alter the disorder or hydrogen bonding capabilities of this region^65^. We selected these positions due to their universal conservation among the ∼500 most closely related PrkC sequences (Fig. 1C) but their greater diversity across a broader evolutionary set (Fig. 1A).

If the conservation of the IDR simply reflects a specific sequence requirement for activation, then both mutations should reduce kinase activity. While the I295N mutation reduced kinase activity ∼4-fold, the A292P increased activity ∼1.25-fold relative to WT (Fig. S4). These findings indicate that the overall conservation of this region cannot be explained exclusively by the need to maximize kinase activity.

We sought to determine whether these IDR mutations are epistatic to the T290A phosphoablative mutation, which could potentially indicate that their primary role is to modulate phosphorylation of Thr 290. Because PrkC^T290A^ exhibits low activity, we measured the combinatorial effects of these mutations by titrating kinase expression. We found that PrkC^T290A^ ^I295N^ showed activity comparable to PrkC^T290A^ at all tested expression levels, suggesting that the negative effect of I295N depends on phosphorylation of Thr 290 (Fig. 4A(ii)). In contrast, PrkC^T290A^ ^A292P^ displayed ∼2-fold higher activity than PrkC^T290A^, indicating that the positive effect of A292P is at least partly independent of Thr 290 phosphorylation (Fig. 4A(iii)). Therefore, the A292P mutation presents a separation-of-function allele for the kinase, partially bypassing the Thr 290 requirement, while increasing total kinase activity. This is consistent with the PrkC^A292P^ variant partially relieving IDR-mediated kinase autoinhibition in a phosphorylation-independent manner.

Interpreting Thr 290 phosphorylation as the uninhibited state and the phosphoablative T290A as the inhibited state, these data suggest that the IDR could function as an inhibitory regulatory domain for kinase activity that prevents subsequent positive feedback. In this context, the A292P mutation increases kinase activity in both states, compromising tight regulation, whereas I295N decreases activity specifically in the uninhibited state. Notably, both mutations reduce the dynamic range from ∼4.6-fold for PrkC^WT^ to ∼2.8-fold and ∼3.0-fold for the I295N and A292P mutants, respectively. This reduction suggests that both mutations compromise the potential role of the IDR as a regulatory domain and could provide a mechanistic explanation for the strong evolutionary conservation in this region.

### PrkC autophosphorylation occurs in trans

In vitro studies have shown that PrkC is capable of autophosphorylation in trans by demonstrating that a catalytic mutant PrkC^K40A^ is phosphorylated when incubated with PrkC^WT23^. However, this result did not determine whether trans-phosphorylation results in active PrkC^K40A^. We sought to determine if trans-activation occurs in vivo and to establish the role, if any, of the IDR in that process. Previous studies and Fig. 2 show that *prkC^K40A^* results in a loss of PrkC-dependent phenotypes in vivo^17,42,66^, but this does not preclude the possibility that the catalytic mutant kinase may retain latent functionality if activated through trans-phosphorylation. Indeed, in some eukaryotic systems, mutation of the conserved catalytic Lys residue does not always completely eliminate kinase function^67,68^.

To test whether PrkC^K40A^ can be functionally activated in trans in vivo, we combined *prkC* chromosomal allelic replacements with the xylose-inducible expression system to titrate expression of a second *prkC* variant (Fig. 4B). This dual-expression strategy allowed us to assess whether PrkC^K40A^ can be trans-phosphorylated and activated by other kinase variants. We performed two complementary experiments in which either *prkC^WT^* was expressed from the native locus while *prkC^K40A^* was titrated from the inducible promoter, or conversely *prkC^K40A^* was expressed from the native locus while *prkC^WT^* was titrated (Fig. 4C). In both approaches, we observed measurable PrkC^K40A^ activity in the presence of PrkC^WT^, indicating functional trans-activation. Our mathematical model (SI Text) estimates that trans-phosphorylated PrkC^K40A^ retains approximately one-third of the catalytic activity of PrkC^WT^. This model illustrates that despite this partial activity, PrkC^K40A^ appears inactive when expressed in isolation due to positive feedback introducing a sharp activation threshold. At the range of expression levels in these experiments, PrkC^K40A^ cannot sustain a population of active kinases, further supporting a model in which intermolecular phosphorylation contributes to PrkC activation in vivo.

### Trans-activation specifically requires IDR phosphorylation of the substrate kinase

To investigate the role of IDR in trans-activation, we focused on two questions: first, whether Thr 290 is required in the activating kinase that catalyzes trans-phosphorylation, and second, whether it is required in the substrate kinase.

To test whether Thr 290 is necessary in the activating kinase, we co-expressed the catalytically inactive mutant *prkC^K40A^* with *prkC^T290A^* (Fig. 4C). As in the case for PrkC^WT^, PrkC^T290A^ activated PrkC^K40A^ in trans. When interpreted through our mathematical model (SI Text), the observed decrease in trans-activation efficiency can be attributed to the generally reduced activity of PrkC^T290A^. This suggests that Thr 290 is not specifically required in the activating kinase to facilitate trans-activation.

We then asked whether the presence of Thr 290 is required in the substrate kinase during trans-activation. To address this question, we used the double mutant PrkC^K40A^ ^T290A^ in which both the catalytic site and the IDR phosphorylation site are disrupted. We titrated its expression alone and in combination with other *prkC* variants, including *prkC^WT^*. In all cases, whether expressed in isolation or co-expressed with any other *prkC* variant, the double mutant showed no detectable activity (Fig. 4A,C). These results demonstrate that phosphorylation of Thr 290 within the IDR is essential in the substrate kinase for its activation in trans.

### Evidence against transcriptional autoregulation of *prkC*

Autoregulation is a common feature of many bacterial signaling systems, where the expression level of a kinase is regulated by the overall activity of the signaling pathway^69,70^. For *prkC*, however, there is currently no direct evidence supporting such a regulatory mechanism. This is partly because *prkC* is embedded in a large operon with multiple reported transcriptional start sites^71^, making regulation difficult to isolate and interpret.

To investigate the possibility of autoregulation of *prkC*, we leveraged our dual-expression system. Consider the case where the fully active *prkC^WT^* is co-expressed with a less active mutant, such as *prkC^T290A^*, at about equivalent levels. If there is no feedback regulation on the native promoter, then the total kinase activity should be identical whether *prkC^WT^* is expressed from the native promoter and *prkC^T290A^* is expressed from the inducible promoter, or vice versa. Conversely, if the native promoter (but not the inducible one) is subject to feedback regulation, then the activity levels in the two scenarios would differ. For example, under positive feedback, the activity would be higher in a strain where *prkC^WT^* is expressed from the native promoter than in a strain where it is expressed from the inducible promoter. Under negative feedback, activity in the former strain would be lower.

In the two cases we tested, pairing *prkC^WT^* with either *prkC^T290A^* or *prkC^K40A^*, we found the kinase activity level measured to be independent of which promoter drove expression of *prkC* (Fig. 4D, S5). These findings strongly argue against feedback regulation acting on the native *prkC* promoter under the conditions tested.

### IDR phosphosite-dependent ligand-induced activation of PrkC

Evidence suggests that the β-lactam antibiotic cefotaxime activates PrkC, most likely by increasing the concentration of its activating ligand. This effect may arise through interference with lipid II processing or another mechanism that perturbs cell wall homeostasis^27,28,72^. Since Thr 290 is important for activation, we next asked whether this site also plays a role in ligand-induced activation in response to cefotaxime.

To measure the role of the IDR Thr 290 during β-lactam treatment, we repeated the kinase co-expression titration experiments with a sub-inhibitory concentration of cefotaxime (Fig. 5A). When PrkC^WT^ was the only kinase variant, cefotaxime resulted in an increase in activation across all levels of expression, up to saturation. In contrast, when only PrkC^T290A^ was present, no cefotaxime-dependent increase in activation was observed, even at expression levels that show robust kinase activity. This suggests that cefotaxime-dependent induction requires Thr 290 phosphorylation.

**Figure 5:**
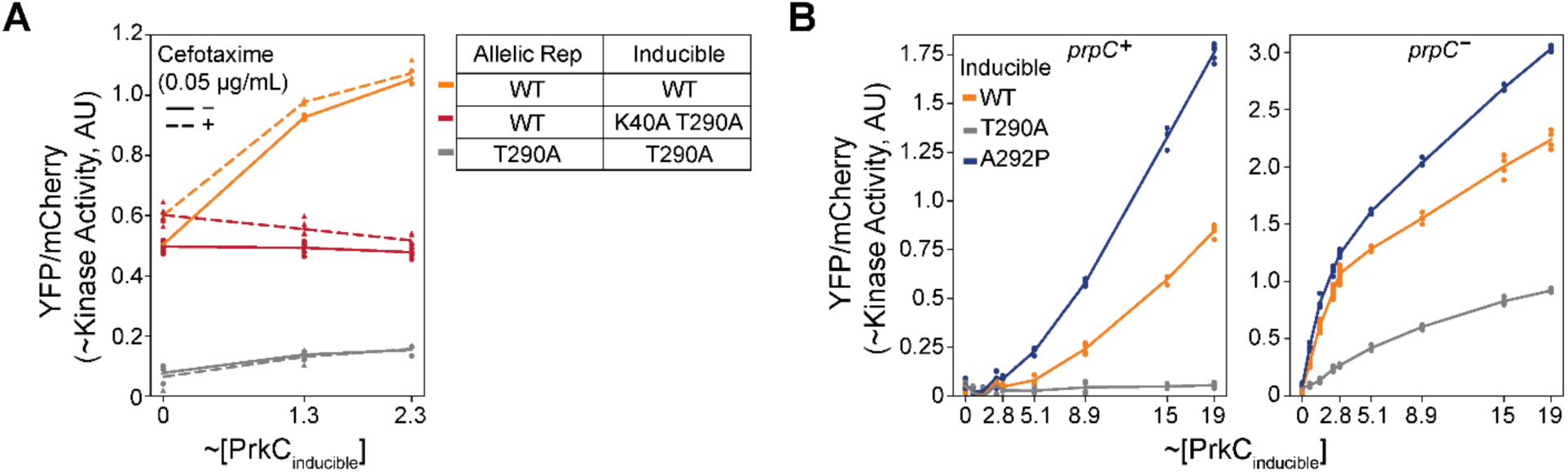
IDR-phosphosite-dependent kinase activation in the presence of cefotaxime or the phosphatase. **A) The effect of Thr 290 on ligand-dependent kinase activation.** Total kinase activity for co-expression strains in the presence (dashed line) or absence (solid line) of sub-MIC concentration cefotaxime (0.05 μg/mL). The addition of cefotaxime increases kinase activation of *prkC^WT^* but not *prkC^T290A^*. Increased expression of *prkC^K40A^ ^T290A^* in the presence of *prkC^WT^* results in a progressive decrease in the ability of cefotaxime to increase kinase activity. **B) IDR-dependent regulation of kinase activation in the presence of the phosphatase.** Dose response of kinase activity in response to titration of *prkC* variants in the presence of (**Left**) or absence of (**Right**) the phosphatase PrpC. In the presence of the phosphatase, *prkC^WT^* display a non-linear threshold behavior, while T290A variants exhibit no increase in kinase activity at up to 19-fold native expression levels. For all panels, each dot represents the mean YFP/mCherry of 3 replicates from biologically independent experiments (n = 3-6 in A, n = 3-16 in B).

The dimerization of PrkC homologs in vitro led to the hypothesis that ligand-induced activation of PrkC occurs through dimerization^19,20^. If this is the case, the failure of PrkC^T290A^ to respond to the ligand may imply that PrkC^T290A^ either fails to form or fails to attain stabilized active dimers. We reasoned that if cefotaxime increases dimer formation or stability in a Thr 290-dependent manner, this effect could be disrupted by high expression of the inactive kinase variant PrkC^K40AT290A^ (Fig. 4B,D) due to molecular interactions between active PrkC^WT^ and inactive PrkC^K40A^ ^T290A^. To test this hypothesis, we co-expressed *prkC^K40A^ ^T290A^* with *prkC^WT^*, with and without cefotaxime (Fig. 5A, red lines). In the absence of cefotaxime, co-expression had no effect on total kinase activity, indicating that PrkC^K40A^ ^T290A^ does not significantly interfere with the function of PrkC^WT^. In the presence of cefotaxime, titration of PrkC^K40A^ ^T290A^ led to a progressive decrease in kinase activity, ultimately approaching the baseline level seen with PrkC^WT^ alone under non-inducing conditions. This suggests that high concentrations of PrkC^K40A^ ^T290A^ effectively block the ligand-mediated increase in kinase activity by creating a dominant pool of kinases incapable of further activation, perhaps by interfering with the formation of dimers with increased activity.

### IDR phosphosite-dependent kinase activation in the presence of the phosphatase

Up to this point, we performed experiments in a phosphatase knockout background (Δ*prpC*) in order to isolate the role of the kinase IDR from the effects of the cognate phosphatase PrpC. In the wildtype background, the PrkC signaling system exhibits very low net kinase activity under these experimental conditions (mid-log growth without antibiotic treatment)^24,50,53^, presumably due to high phosphatase activity. Prior studies demonstrated that sufficient PrkC overexpression, even in the presence of the phosphatase, yields measurable changes in downstream gene expression. Therefore, to measure the effect of IDR variants on kinase activation in the presence of the phosphatase, we extended our overexpression system to ∼19-fold the native levels of PrkC (Fig. 5B, S3).

In the presence of the phosphatase, expression of either *prkC^WT^* or the highly active *prkC^A292P^*, which showed no detectable threshold in the absence of the phosphatase, resulted in a threshold-like behavior of kinase activity, reflecting the effect of the autoactivation feedback. These results are consistent with the phosphatase reducing the net activity of individual kinases, thereby shifting the activation threshold (SI Text). Titrating expression of *prkC^T290A^* resulted in no detectable increase in activity even at high expression levels in the *prpC*^+^ background. This suggests that the T290A mutation results in a higher activation threshold, compared to the *prpC^-^* background, that cannot be crossed even at high levels of kinase expression.

## Discussion

One of the enduring open questions in bacterial signaling has been how prokaryotic Hanks-type Ser/Thr kinase-phosphatase signaling pathways activate. Although understudied, this class of signaling systems is widespread throughout prokaryotes and is well recognized to regulate important processes such as cell growth, antibiotic tolerance, and virulence.

In this study, we found that many of these kinases have an IDR adjacent to the kinase domain. In addition to the conservation of this region, a potential phosphothreonine or phosphoserine is typically found ∼20 residues from the kinase domain. The consistent location of this phosphosite within the IDR suggests a physical constraint on its placement. We showed that a phosphoablative mutation at Thr 290 in the kinase PrkC from *B. subtilis* strongly decreases kinase activity in vivo, corroborated by phenotypic changes, downstream gene expression, and a synthetic reporter of kinase activity. We found that IDR variants affect kinase activation in a phosphosite-dependent manner, that PrkC autoactivation can occur in trans, and that the phosphoablative T290A IDR mutation impairs response to cefotaxime. We showed that at sufficiently high levels of kinase expression, positive feedback due to autoactivation results in sustained pathway activity, whereas in the wildtype background, the physiological concentration of PrkC is not high enough resulting in low activity.

It has long been hypothesized that PrkC and its homologs may dimerize in response to ligand binding, but there is currently a lack of direct evidence for this occurring in vivo. Our results in a kinase co-expression system with the addition of cefotaxime are consistent with ligand binding-mediated dimerization leading to kinase activation in a Thr 290-dependent manner. However, obtaining more direct evidence will require a more complete understanding of the ligand and its binding to the receptor.

Taken together, our data suggest a possible model (Fig. 6) in which the IDR functions as a phosphorylation-dependent cis-inhibitory domain that regulates both trans-activation and ligand binding-mediated signaling. This dual-layer regulatory architecture suggests that the IDR can function as a sophisticated regulatory element, as seen in analogous systems with similar architectures. If this is the case, the IDR may integrate the response of the signaling pathway to additional cofactors such as other proteins, compounds, or even interactions with the phosphatase.

**Figure 6:**
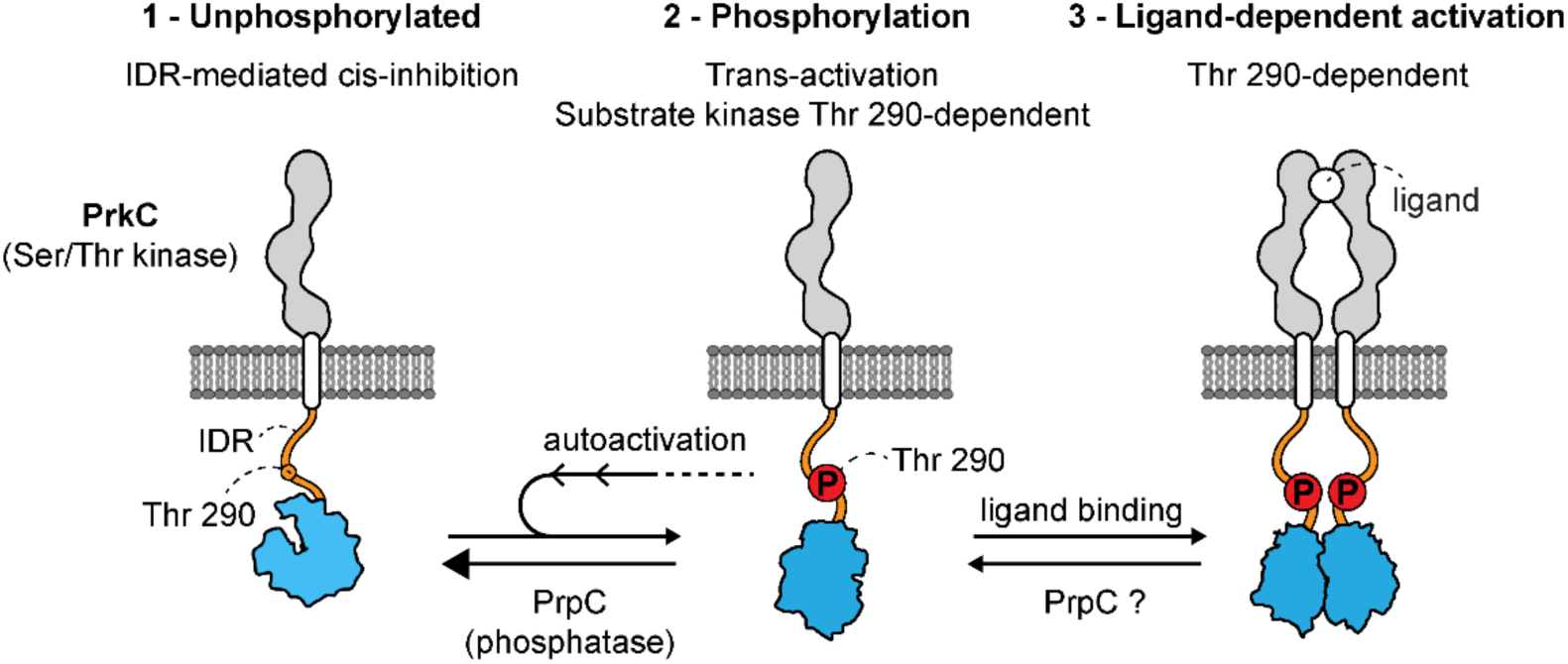
Model for IDR phosphorylation-dependent activation of the bacterial Hanks-type Ser/Thr kinase PrkC. Prior to IDR phosphorylation at Thr 290, PrkC is in an inactive state, i.e. cis-inhibited. Phosphorylation of the IDR relieves cis-inhibition, enabling IDR-dependent trans-activation by another PrkC kinase and subsequent trans-phosphorylation of other PrkC kinases, resulting in positive feedback. The phosphatase PrpC can likely dephosphorylate monomeric and dimeric PrkC. Ligand binding-mediated kinase activation is also Thr 290 phosphorylation-dependent, and likely involves PrkC dimerization. Activated PrkC results in downstream signaling, including changes in gene expression and physiological phenotypes.

Precisely how the IDR regulates cis-autoinhibition remains unclear due to its unstructured nature. In addition to the Thr 290 site, our IDR variant data (Fig. 4A) indicate that surrounding residues may have a function beyond modulating phosphorylation efficiency, potentially through physical interactions with the catalytic domain. In the unphosphorylated state, the IDR may inhibit activation either by interacting with the kinase active site or by preventing phosphorylation of the activation loop. Evidence from eukaryotic Hanks-type Ser/Thr kinases, where kinase trans-activation of a catalytic mutant requires phosphorylation of the activation loop^64,68,73^, suggests that in PrkC^T290A^ the activation loop could be blocked in the substrate kinase. Phosphorylation of the IDR likely relieves this inhibition, possibly by promoting release of the IDR from the kinase domain, enabling full kinase activation and dimerization. Importantly, IDR phosphorylation appears to be specifically required in the substrate kinase during trans-activation, supporting its role as a cis-autoinhibitory domain that regulates PrkC activity. The PrkC^A292P^ variant further supports this model, as the A292P separation-of-function mutation compromises the inhibitory role of the IDR while still allowing for phosphorylation responsiveness. We note, however, that it is also possible that IDR phosphorylation directly promotes kinase activation, rather than relieving inhibition, though the physical mechanism behind the requirement of Thr 290 phosphorylation in activating K40A variants (Fig. 4C) is unclear.

Autophosphorylation in trans is a common activation mechanism for bacterial kinases^74^. For PrkC, this autophosphorylation step, in addition to cis-inhibition, could provide a barrier to spurious kinase activation and the subsequent strong positive feedback, thereby filtering out noise^62,75^.

Intrinsically disordered regions have been implicated in generating subcellular organization in diverse species and systems^76–80^. Although less common in bacteria, there are several examples in which disordered domains regulate protein-protein interactions, sensing of a signal, or the formation of phase condensates in vitro or in vivo^81–90^. In prokaryotes, the role of IDRs in the subcellular organization of Hanks-type Ser/Thr kinases or their substrates remains unclear. However, subcellular organization of PrkC was previously observed in *B. subtilis*^91^, and a recent study in *S. pneumoniae* suggests its kinase juxtamembrane IDR may be involved in organization and recruitment of divisome proteins^27^.

Our results demonstrate the critical role that kinase IDR phosphorylation plays in antibiotic sensitivity (Fig. 2). In *B. subtilis*, PrkC-dependent changes in antibiotic sensitivity have been previously observed, including to cefotaxime, corroborated by microarray^48,92^ and our synthetic kinase activity reporter^24^. Beyond *B. subtilis*, PASTA kinases have been broadly implicated in β-lactam sensitivity in clinically important pathogens, including in *E. faecalis*^43,93–95^, *S. aureus*^96^, and *S. pneumoniae*^28,97,98^. This suggests that the IDR-mediated mechanism of activation described here may present a conserved target for antimicrobial development.

Together, our results support an IDR phosphorylation-dependent activation mechanism for the bacterial Hanks-type Ser/Thr kinase PrkC. These findings expand our fundamental understanding of a widespread class of bacterial signaling systems and demonstrate that phosphorylation of bacterial IDRs can have important regulatory and phenotypic consequences. Given the well-established role of PASTA kinases in growth, virulence, and antibiotic sensitivity, our work highlights the kinase juxtamembrane IDR as a potential target for antimicrobial development. Future work will be critical for understanding the potential regulatory role of the IDR in signal integration, its cell biological consequences, and the broader conservation of this mechanism across prokaryotes.

## Author Contributions

All authors conceived the project and analyzed data. N.G. performed experiments and E.A.L and E.L. provided mentorship and supervision. E.L contributed a mathematical model, and N.G. and E.L. performed computational and mathematical analyses. All authors wrote, reviewed, and edited the manuscript. E.A.L. acquired resources and E.A.L and E.L. acquired funding.

## Data availability

All relevant data are within the manuscript and its Supporting Information files.

## Acknowledgements

We thank Jonathan Dworkin and Mark Goulian for generously providing strains and plasmids. We thank Jonathan Dworkin for helpful advice and comments on the manuscript. This work was funded by NIH grant R35GM147429 to EAL and by NSF grant MCB1946944 to EL.

## Conflict of Interest

The authors declare no conflict of interest.

## Notes

### Competing Interest Statement

The authors have declared no competing interest.

## References

1 Hanks, S. K., Quinn, A. M. & Hunter, T. The protein kinase family: conserved features and deduced phylogeny of the catalytic domains. Science 241, 42–52 (1988). 10.1126/science.3291115

2 Munoz-Dorado, J., Inouye, S. & Inouye, M. A gene encoding a protein serine/threonine kinase is required for normal development of M. xanthus, a gram-negative bacterium. Cell 67, 995–1006 (1991). 10.1016/0092-8674(91)90372-6

3 Stancik, I. A. et al. Serine/Threonine Protein Kinases from Bacteria, Archaea and Eukarya Share a Common Evolutionary Origin Deeply Rooted in the Tree of Life. J Mol Biol 430, 27–32 (2018). 10.1016/j.jmb.2017.11.004

4 O’Boyle, B. et al. An atlas of bacterial serine-threonine kinases reveals functional diversity and key distinctions from eukaryotic kinases. Sci Signal 18, eadt8686 (2025). 10.1126/scisignal.adt8686

5 Pereira, S. F., Goss, L. & Dworkin, J. Eukaryote-like serine/threonine kinases and phosphatases in bacteria. Microbiol Mol Biol Rev 75, 192–212 (2011). 10.1128/MMBR.00042-10

6 Dworkin, J. Ser/Thr phosphorylation as a regulatory mechanism in bacteria. Curr Opin Microbiol 24, 47–52 (2015). 10.1016/j.mib.2015.01.005

7 Grunfeld, N., Levine, E. & Libby, E. Experimental measurement and computational prediction of bacterial Hanks-type Ser/Thr signaling system regulatory targets. Mol Microbiol 122, 152–164 (2024). 10.1111/mmi.15220

8 Yeats, C., Finn, R. D. & Bateman, A. The PASTA domain: a beta-lactam-binding domain. Trends Biochem Sci 27, 438 (2002). 10.1016/s0968-0004(02)02164-3

9 Pensinger, D. A., Schaenzer, A. J. & Sauer, J. D. Do Shoot the Messenger: PASTA Kinases as Virulence Determinants and Antibiotic Targets. Trends Microbiol 26, 56–69 (2018). 10.1016/j.tim.2017.06.010

10 Squeglia, F. et al. Chemical basis of peptidoglycan discrimination by PrkC, a key kinase involved in bacterial resuscitation from dormancy. J Am Chem Soc 133, 20676–20679 (2011). 10.1021/ja208080r

11 Huemer, M. et al. Serine-threonine phosphoregulation by PknB and Stp contributes to quiescence and antibiotic tolerance in Staphylococcus aureus. Sci Signal 16, eabj8194 (2023). 10.1126/scisignal.abj8194

12 Vornhagen, J. et al. Kinase Inhibitors that Increase the Sensitivity of Methicillin Resistant Staphylococcus aureus to beta-Lactam Antibiotics. Pathogens 4, 708–721 (2015). 10.3390/pathogens4040708

13 King, A. & Blackledge, M. S. Evaluation of small molecule kinase inhibitors as novel antimicrobial and antibiofilm agents. Chem Biol Drug Des 98, 1038–1064 (2021). 10.1111/cbdd.13962

14 Konaklieva, M. I. & Plotkin, B. J. Utilization of Existing Human Kinase Inhibitors as Scaffolds in the Development of New Antimicrobials. Antibiotics (Basel) 12 (2023). 10.3390/antibiotics12091418

15 Nagarajan, S. N., Lenoir, C. & Grangeasse, C. Recent advances in bacterial signaling by serine/threonine protein kinases. Trends Microbiol 30, 553–566 (2022). 10.1016/j.tim.2021.11.005

16 Hardt, P. et al. The cell wall precursor lipid II acts as a molecular signal for the Ser/Thr kinase PknB of Staphylococcus aureus. Int J Med Microbiol 307, 1–10 (2017). 10.1016/j.ijmm.2016.12.001

17 Shah, I. M., Laaberki, M. H., Popham, D. L. & Dworkin, J. A eukaryotic-like Ser/Thr kinase signals bacteria to exit dormancy in response to peptidoglycan fragments. Cell 135, 486–496 (2008). 10.1016/j.cell.2008.08.039

18 Shah, I. M. & Dworkin, J. Induction and regulation of a secreted peptidoglycan hydrolase by a membrane Ser/Thr kinase that detects muropeptides. Mol Microbiol 75, 1232–1243 (2010). 10.1111/j.1365-2958.2010.07046.x

19 Barthe, P., Mukamolova, G. V., Roumestand, C. & Cohen-Gonsaud, M. The structure of PknB extracellular PASTA domain from mycobacterium tuberculosis suggests a ligand-dependent kinase activation. Structure 18, 606–615 (2010). 10.1016/j.str.2010.02.013

20 Labbe, B. D. & Kristich, C. J. Growth- and Stress-Induced PASTA Kinase Phosphorylation in Enterococcus faecalis. J Bacteriol 199 (2017). 10.1128/JB.00363-17

21 Madec, E. et al. Mass spectrometry and site-directed mutagenesis identify several autophosphorylated residues required for the activity of PrkC, a Ser/Thr kinase from Bacillus subtilis. J Mol Biol 330, 459–472 (2003). 10.1016/s0022-2836(03)00579-5

22 Gupta, M., Sajid, A., Arora, G., Tandon, V. & Singh, Y. Forkhead-associated domain-containing protein Rv0019c and polyketide-associated protein PapA5, from substrates of serine/threonine protein kinase PknB to interacting proteins of Mycobacterium tuberculosis. J Biol Chem 284, 34723–34734 (2009). 10.1074/jbc.M109.058834

23 Shi, L. et al. Cross-phosphorylation of bacterial serine/threonine and tyrosine protein kinases on key regulatory residues. Front Microbiol 5, 495 (2014). 10.3389/fmicb.2014.00495

24 Zheng, C. R., Singh, A., Libby, A., Silver, P. A. & Libby, E. A. Modular and Single-Cell Sensors of Bacterial Ser/Thr Kinase Activity. ACS Synth Biol 10, 2340–2350 (2021). 10.1021/acssynbio.1c00250

25 Manuse, S., Fleurie, A., Zucchini, L., Lesterlin, C. & Grangeasse, C. Role of eukaryotic-like serine/threonine kinases in bacterial cell division and morphogenesis. FEMS Microbiol Rev 40, 41–56 (2016). 10.1093/femsre/fuv041

26 Duran, R. et al. Conserved autophosphorylation pattern in activation loops and juxtamembrane regions of Mycobacterium tuberculosis Ser/Thr protein kinases. Biochem Biophys Res Commun 333, 858–867 (2005). 10.1016/j.bbrc.2005.05.173

27 Hamidi, M. et al. The juxtamembrane domain of StkP is phosphorylated and influences cell division in Streptococcus pneumoniae. mBio 16, e0379924 (2025). 10.1128/mbio.03799-24

28 Ulrych, A. et al. Cell Wall Stress Stimulates the Activity of the Protein Kinase StkP of Streptococcus pneumoniae, Leading to Multiple Phosphorylation. J Mol Biol 433, 167319 (2021). 10.1016/j.jmb.2021.167319

29 Macek, B. et al. The serine/threonine/tyrosine phosphoproteome of the model bacterium Bacillus subtilis. Mol Cell Proteomics 6, 697–707 (2007). 10.1074/mcp.M600464-MCP200

30 Ravikumar, V. et al. Quantitative phosphoproteome analysis of Bacillus subtilis reveals novel substrates of the kinase PrkC and phosphatase PrpC. Mol Cell Proteomics 13, 1965–1978 (2014). 10.1074/mcp.M113.035949

31 Ward, J. J., Sodhi, J. S., McGuffin, L. J., Buxton, B. F. & Jones, D. T. Prediction and functional analysis of native disorder in proteins from the three kingdoms of life. J Mol Biol 337, 635–645 (2004). 10.1016/j.jmb.2004.02.002

32 Iakoucheva, L. M. et al. The importance of intrinsic disorder for protein phosphorylation. Nucleic Acids Res 32, 1037–1049 (2004). 10.1093/nar/gkh253

33 van der Lee, R. et al. Classification of intrinsically disordered regions and proteins. Chem Rev 114, 6589–6631 (2014). 10.1021/cr400525m

34 Oldfield, C. J. & Dunker, A. K. Intrinsically disordered proteins and intrinsically disordered protein regions. Annu Rev Biochem 83, 553–584 (2014). 10.1146/annurev-biochem-072711-164947

35 Jumper, J. et al. Highly accurate protein structure prediction with AlphaFold. Nature 596, 583–589 (2021). 10.1038/s41586-021-03819-2

36 Abramson, J. et al. Accurate structure prediction of biomolecular interactions with AlphaFold 3. Nature 630, 493–500 (2024). 10.1038/s41586-024-07487-w

37 Kumar, S., Stecher, G., Suleski, M. & Hedges, S. B. TimeTree: A Resource for Timelines, Timetrees, and Divergence Times. Mol Biol Evol 34, 1812–1819 (2017). 10.1093/molbev/msx116

38 Macek, B. et al. Phosphoproteome analysis of E. coli reveals evolutionary conservation of bacterial Ser/Thr/Tyr phosphorylation. Mol Cell Proteomics 7, 299–307 (2008). 10.1074/mcp.M700311-MCP200

39 Prisic, S. et al. Extensive phosphorylation with overlapping specificity by Mycobacterium tuberculosis serine/threonine protein kinases. Proc Natl Acad Sci U S A 107, 7521–7526 (2010). 10.1073/pnas.0913482107

40 von Heijne, G. Membrane protein structure prediction. Hydrophobicity analysis and the positive-inside rule. J Mol Biol 225, 487–494 (1992). 10.1016/0022-2836(92)90934-c

41 Gaidenko, T. A., Kim, T. J. & Price, C. W. The PrpC serine-threonine phosphatase and PrkC kinase have opposing physiological roles in stationary-phase Bacillus subtilis cells. J Bacteriol 184, 6109–6114 (2002). 10.1128/JB.184.22.6109-6114.2002

42 Madec, E., Laszkiewicz, A., Iwanicki, A., Obuchowski, M. & Seror, S. Characterization of a membrane-linked Ser/Thr protein kinase in Bacillus subtilis, implicated in developmental processes. Mol Microbiol 46, 571–586 (2002). 10.1046/j.1365-2958.2002.03178.x

43 Kristich, C. J., Wells, C. L. & Dunny, G. M. A eukaryotic-type Ser/Thr kinase in Enterococcus faecalis mediates antimicrobial resistance and intestinal persistence. Proc Natl Acad Sci U S A 104, 3508–3513 (2007). 10.1073/pnas.0608742104

44 Cuenot, E. et al. The Ser/Thr Kinase PrkC Participates in Cell Wall Homeostasis and Antimicrobial Resistance in Clostridium difficile. Infect Immun 87 (2019). 10.1128/IAI.00005-19

45 Beltramini, A. M., Mukhopadhyay, C. D. & Pancholi, V. Modulation of cell wall structure and antimicrobial susceptibility by a Staphylococcus aureus eukaryote-like serine/threonine kinase and phosphatase. Infect Immun 77, 1406–1416 (2009). 10.1128/IAI.01499-08

46 Pancholi, V., Boel, G. & Jin, H. Streptococcus pyogenes Ser/Thr kinase-regulated cell wall hydrolase is a cell division plane-recognizing and chain-forming virulence factor. J Biol Chem 285, 30861–30874 (2010). 10.1074/jbc.M110.153825

47 Bisicchia, P. et al. The essential YycFG two-component system controls cell wall metabolism in Bacillus subtilis. Mol Microbiol 65, 180–200 (2007). 10.1111/j.1365-2958.2007.05782.x

48 Dubrac, S., Bisicchia, P., Devine, K. M. & Msadek, T. A matter of life and death: cell wall homeostasis and the WalKR (YycGF) essential signal transduction pathway. Mol Microbiol 70, 1307–1322 (2008). 10.1111/j.1365-2958.2008.06483.x

49 Salzberg, L. I. et al. The WalRK (YycFG) and sigma(I) RsgI regulators cooperate to control CwlO and LytE expression in exponentially growing and stressed Bacillus subtilis cells. Mol Microbiol 87, 180–195 (2013). 10.1111/mmi.12092

50 Libby, E. A., Goss, L. A. & Dworkin, J. The Eukaryotic-Like Ser/Thr Kinase PrkC Regulates the Essential WalRK Two-Component System in Bacillus subtilis. PLoS Genet 11, e1005275 (2015). 10.1371/journal.pgen.1005275

51 Kobayashi, K. et al. Identification and characterization of a novel polysaccharide deacetylase C (PdaC) from Bacillus subtilis. J Biol Chem 287, 9765–9776 (2012). 10.1074/jbc.M111.329490

52 Yamamoto, H. et al. Post-translational control of vegetative cell separation enzymes through a direct interaction with specific inhibitor IseA in Bacillus subtilis. Mol Microbiol 70, 168–182 (2008). 10.1111/j.1365-2958.2008.06398.x

53 Libby, E. A., Reuveni, S. & Dworkin, J. Multisite phosphorylation drives phenotypic variation in (p)ppGpp synthetase-dependent antibiotic tolerance. Nat Commun 10, 5133 (2019). 10.1038/s41467-019-13127-z

54 Nolen, B., Taylor, S. & Ghosh, G. Regulation of protein kinases; controlling activity through activation segment conformation. Mol Cell 15, 661–675 (2004). 10.1016/j.molcel.2004.08.024

55 Endicott, J. A., Noble, M. E. & Johnson, L. N. The structural basis for control of eukaryotic protein kinases. Annu Rev Biochem 81, 587–613 (2012). 10.1146/annurev-biochem-052410-090317

56 Batchelor, E. & Goulian, M. Robustness and the cycle of phosphorylation and dephosphorylation in a two-component regulatory system. Proc Natl Acad Sci U S A 100, 691–696 (2003). 10.1073/pnas.0234782100

57 Miyashiro, T. & Goulian, M. Stimulus-dependent differential regulation in the Escherichia coli PhoQ PhoP system. Proc Natl Acad Sci U S A 104, 16305–16310 (2007). 10.1073/pnas.0700025104

58 Gao, R. & Stock, A. M. Quantitative Kinetic Analyses of Shutting Off a Two-Component System. mBio 8 (2017). 10.1128/mBio.00412-17

59 Salazar, M. E., Podgornaia, A. I. & Laub, M. T. The small membrane protein MgrB regulates PhoQ bifunctionality to control PhoP target gene expression dynamics. Mol Microbiol 102, 430–445 (2016). 10.1111/mmi.13471

60 Butcher, R. J. & Tabor, J. J. Real-time detection of response regulator phosphorylation dynamics in live bacteria. Proc Natl Acad Sci U S A 119, e2201204119 (2022). 10.1073/pnas.2201204119

61 Rao, S. D. & Igoshin, O. A. Overlaid positive and negative feedback loops shape dynamical properties of PhoPQ two-component system. PLoS Comput Biol 17, e1008130 (2021). 10.1371/journal.pcbi.1008130

62 Kollmann, M., Lovdok, L., Bartholome, K., Timmer, J. & Sourjik, V. Design principles of a bacterial signalling network. Nature 438, 504–507 (2005). 10.1038/nature04228

63 Lovdok, L. et al. Role of translational coupling in robustness of bacterial chemotaxis pathway. PLoS Biol 7, e1000171 (2009). 10.1371/journal.pbio.1000171

64 Reinhardt, R. & Leonard, T. A. A critical evaluation of protein kinase regulation by activation loop autophosphorylation. Elife 12 (2023). 10.7554/eLife.88210

65 Radivojac, P. et al. Intrinsic disorder and functional proteomics. Biophys J 92, 1439–1456 (2007). 10.1529/biophysj.106.094045

66 Absalon, C. et al. CpgA, EF-Tu and the stressosome protein YezB are substrates of the Ser/Thr kinase/phosphatase couple, PrkC/PrpC, in Bacillus subtilis. Microbiology (Reading) 155, 932–943 (2009). 10.1099/mic.0.022475-0

67 Iyer, G. H., Garrod, S., Woods, V. L., Jr. & Taylor, S. S. Catalytic independent functions of a protein kinase as revealed by a kinase-dead mutant: study of the Lys72His mutant of cAMP-dependent kinase. J Mol Biol 351, 1110–1122 (2005). 10.1016/j.jmb.2005.06.011

68 Eyers, P. A. et al. Regulation of the G(2)/M transition in Xenopus oocytes by the cAMP-dependent protein kinase. J Biol Chem 280, 24339–24346 (2005). 10.1074/jbc.M412442200

69 Goulian, M. Two-component signaling circuit structure and properties. Curr Opin Microbiol 13, 184–189 (2010). 10.1016/j.mib.2010.01.009

70 Groisman, E. A. Feedback Control of Two-Component Regulatory Systems. Annu Rev Microbiol 70, 103–124 (2016). 10.1146/annurev-micro-102215-095331

71 Nicolas, P. et al. Condition-dependent transcriptome reveals high-level regulatory architecture in Bacillus subtilis. Science 335, 1103–1106 (2012). 10.1126/science.1206848

72 Maestro, B. et al. Recognition of peptidoglycan and beta-lactam antibiotics by the extracellular domain of the Ser/Thr protein kinase StkP from Streptococcus pneumoniae. FEBS Lett 585, 357–363 (2011). 10.1016/j.febslet.2010.12.016

73 Haydon, C. E. et al. Identification of novel phosphorylation sites on Xenopus laevis Aurora A and analysis of phosphopeptide enrichment by immobilized metal-affinity chromatography. Mol Cell Proteomics 2, 1055–1067 (2003). 10.1074/mcp.M300054-MCP200

74 Zschiedrich, C. P., Keidel, V. & Szurmant, H. Molecular Mechanisms of Two-Component Signal Transduction. J Mol Biol 428, 3752–3775 (2016). 10.1016/j.jmb.2016.08.003

75 Mascher, T., Helmann, J. D. & Unden, G. Stimulus perception in bacterial signal-transducing histidine kinases. Microbiol Mol Biol Rev 70, 910–938 (2006). 10.1128/MMBR.00020-06

76 Banani, S. F., Lee, H. O., Hyman, A. A. & Rosen, M. K. Biomolecular condensates: organizers of cellular biochemistry. Nat Rev Mol Cell Biol 18, 285–298 (2017). 10.1038/nrm.2017.7

77 Holehouse, A. S. & Kragelund, B. B. The molecular basis for cellular function of intrinsically disordered protein regions. Nat Rev Mol Cell Biol 25, 187–211 (2024). 10.1038/s41580-023-00673-0

78 Wright, P. E. & Dyson, H. J. Intrinsically disordered proteins in cellular signalling and regulation. Nat Rev Mol Cell Biol 16, 18–29 (2015). 10.1038/nrm3920

79 Martin, E. W. & Holehouse, A. S. Intrinsically disordered protein regions and phase separation: sequence determinants of assembly or lack thereof. Emerg Top Life Sci 4, 307–329 (2020). 10.1042/ETLS20190164

80 Holehouse, A. S. & Pappu, R. V. Functional Implications of Intracellular Phase Transitions. Biochemistry 57, 2415–2423 (2018). 10.1021/acs.biochem.7b01136

81 Azaldegui, C. A., Vecchiarelli, A. G. & Biteen, J. S. The emergence of phase separation as an organizing principle in bacteria. Biophys J 120, 1123–1138 (2021). 10.1016/j.bpj.2020.09.023

82 Buske, P. J., Mittal, A., Pappu, R. V. & Levin, P. A. An intrinsically disordered linker plays a critical role in bacterial cell division. Semin Cell Dev Biol 37, 3–10 (2015). 10.1016/j.semcdb.2014.09.017

83 Brunet, Y. R., Habib, C., Brogan, A. P., Artzi, L. & Rudner, D. Z. Intrinsically disordered protein regions are required for cell wall homeostasis in Bacillus subtilis. Genes Dev 36, 970–984 (2022). 10.1101/gad.349895.122

84 Cohan, M. C., Eddelbuettel, A. M. P., Levin, P. A. & Pappu, R. V. Dissecting the Functional Contributions of the Intrinsically Disordered C-terminal Tail of Bacillus subtilis FtsZ. J Mol Biol 432, 3205–3221 (2020). 10.1016/j.jmb.2020.03.008

85 Gardner, K. A., Moore, D. A. & Erickson, H. P. The C-terminal linker of Escherichia coli FtsZ functions as an intrinsically disordered peptide. Mol Microbiol 89, 264–275 (2013). 10.1111/mmi.12279

86 Holmes, J. A. et al. Caulobacter PopZ forms an intrinsically disordered hub in organizing bacterial cell poles. Proc Natl Acad Sci U S A 113, 12490–12495 (2016). 10.1073/pnas.1602380113

87 Sasazawa, M., Tomares, D. T., Childers, W. S. & Saurabh, S. Biomolecular condensates as stress sensors and modulators of bacterial signaling. PLoS Pathog 20, e1012413 (2024). 10.1371/journal.ppat.1012413

88 Saurabh, S. et al. ATP-responsive biomolecular condensates tune bacterial kinase signaling. Sci Adv 8, eabm6570 (2022). 10.1126/sciadv.abm6570

89 Lasker, K. et al. The material properties of a bacterial-derived biomolecular condensate tune biological function in natural and synthetic systems. Nat Commun 13, 5643 (2022). 10.1038/s41467-022-33221-z

90 Nandana, V. et al. The BR-body proteome contains a complex network of protein-protein and protein-RNA interactions. Cell Rep 42, 113229 (2023). 10.1016/j.celrep.2023.113229

91 Pompeo, F., Byrne, D., Mengin-Lecreulx, D. & Galinier, A. Dual regulation of activity and intracellular localization of the PASTA kinase PrkC during Bacillus subtilis growth. Sci Rep 8, 1660 (2018). 10.1038/s41598-018-20145-2

92 Hutter, B. & John, G. T. Evaluation of OxoPlate for real-time assessment of antibacterial activities. Curr Microbiol 48, 57–61 (2004). 10.1007/s00284-003-4095-4

93 Iannetta, A. A. et al. IreK-Mediated, Cell Wall-Protective Phosphorylation in Enterococcus faecalis. J Proteome Res 20, 5131–5144 (2021). 10.1021/acs.jproteome.1c00635

94 VanZeeland, N. E., Schultz, K. M., Klug, C. S. & Kristich, C. J. Multisite Phosphorylation Regulates GpsB Function in Cephalosporin Resistance of Enterococcus faecalis. J Mol Biol 435, 168216 (2023). 10.1016/j.jmb.2023.168216

95 Gilmore, M. S. et al. Genes Contributing to the Unique Biology and Intrinsic Antibiotic Resistance of Enterococcus faecalis. mBio 11 (2020). 10.1128/mBio.02962-20

96 Chatterjee, A., Poon, R. & Chatterjee, S. S. Stp1 Loss of Function Promotes beta-Lactam Resistance in Staphylococcus aureus That Is Independent of Classical Genes. Antimicrob Agents Chemother 64 (2020). 10.1128/AAC.02222-19

97 Huang, Y. Y. et al. Sublethal beta-lactam antibiotics induce PhpP phosphatase expression and StkP kinase phosphorylation in PBP-independent beta-lactam antibiotic resistance of Streptococcus pneumoniae. Biochem Biophys Res Commun 503, 2000–2008 (2018). 10.1016/j.bbrc.2018.07.148

98 Tamber, S., Schwartzman, J. & Cheung, A. L. Role of PknB kinase in antibiotic resistance and virulence in community-acquired methicillin-resistant Staphylococcus aureus strain USA300. Infect Immun 78, 3637–3646 (2010). 10.1128/IAI.00296-10

