## Supplemental Information for "Phosphorylation of a conserved intrinsically disordered region is necessary for activation of a bacterial Hanks-type Ser/Thr kinase signaling pathway"

**Table of Contents**


Supplementary Figures …… 3-7

Supplementary Text …… 8-13

Materials & Methods …… 14-17

Supplementary Tables …… 18-28

Supplementary Information References …… 29

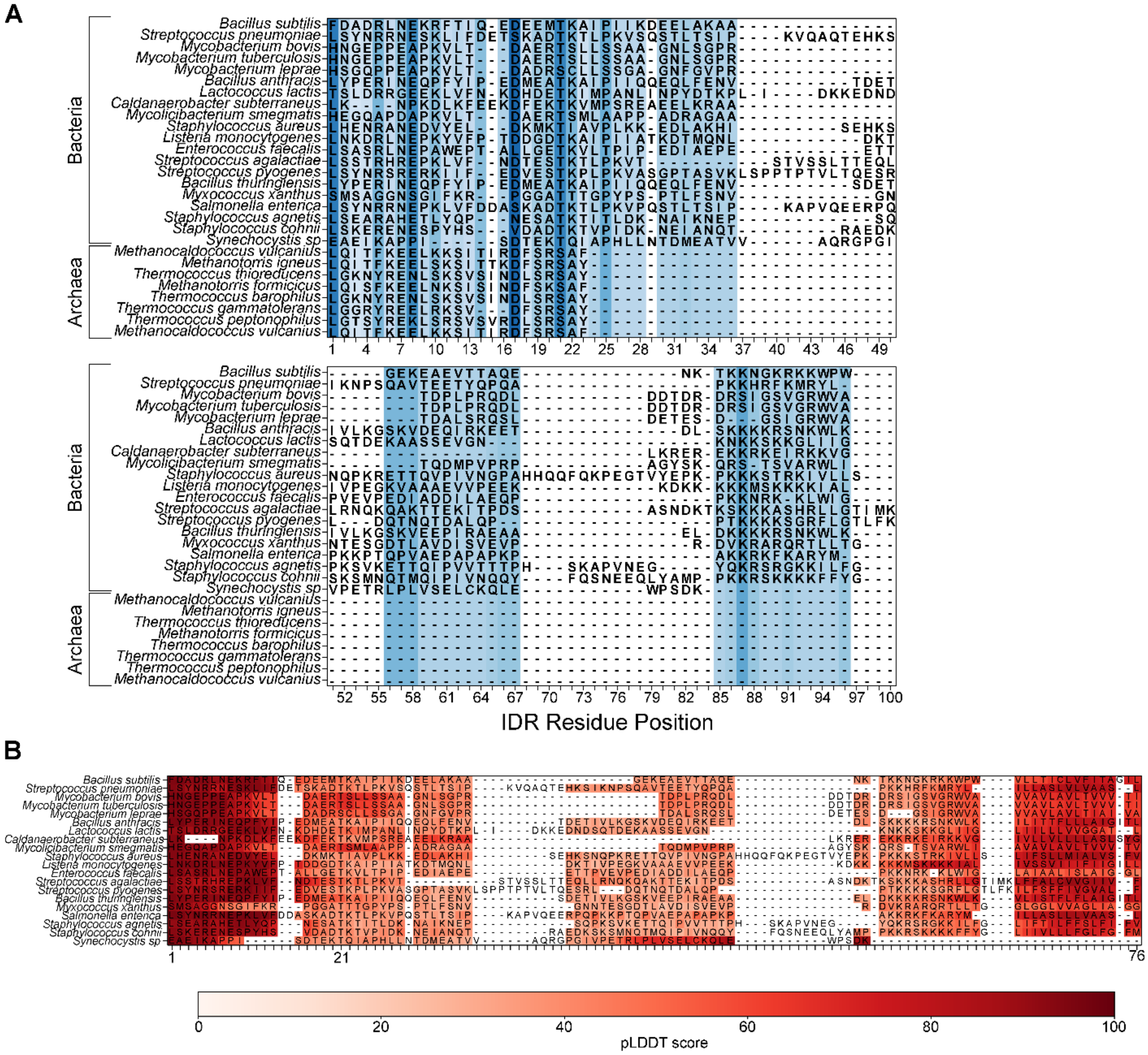


**Figure S1: Conservation of the kinase domain-proximal IDR in prokaryotic Ser/Thr kinases.**

A) Multiple sequence alignment of prokaryotic Hanks-type Ser/Thr kinase IDRs in bacterial and archaeal species, shown relative to PrkC from *B. subtilis* (top row). Shading highlights the most highly conserved positions (dark blue) and variable residues (light blue to white). Position 21 corresponds to *B. subtilis* Thr 290.

B) Per-residue disorder scores mapped onto aligned bacterial kinases from A. Residues are colored by prediction confidence (pLDDT from AlphaFold: dark red = high confidence/low disorder, light = low confidence/high disorder).

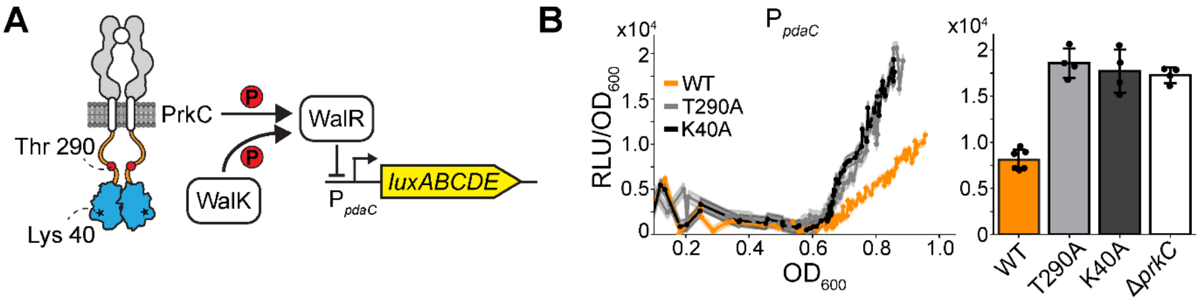


**Figure S2: A native WalR-regulon transcriptional reporter shows PrkC Thr 290 phosphosite dependent regulation.**

A) **Schematic of a WalR-based transcriptional reporter for PrkC activity.** PrkC and WalK phosphorylate WalR, increasing repression of *pdaC* in stationary phase, reflected in activity of $P_{pdaC}$-*lux*.

B) **A *pdaC* reporter shows Thr 290 phosphosite-dependent regulation.** Normalized luminescence (RLU/$OD_{600}$) as a function of growth ($OD_{600}$). **Left:** Cultures of *prkC*$^{WT}$ (orange), *prkC*$^{T290A}$ (grey), and *prkC*$^{K40A}$ (black) allelic replacement strains were grown in microplates from early log into stationary phase. Dots represent means of 3 replicates, shading the standard deviation. ***Right***: Bar plot show means at early stationary phase. Error bars indicate the standard error from ≥3 independent experiments (dots), including the representative experiment shown on the left. Both *prkC*$^{T290A}$ and *prkC*$^{K40A}$ mutants result in a ~2-fold increase in signal.

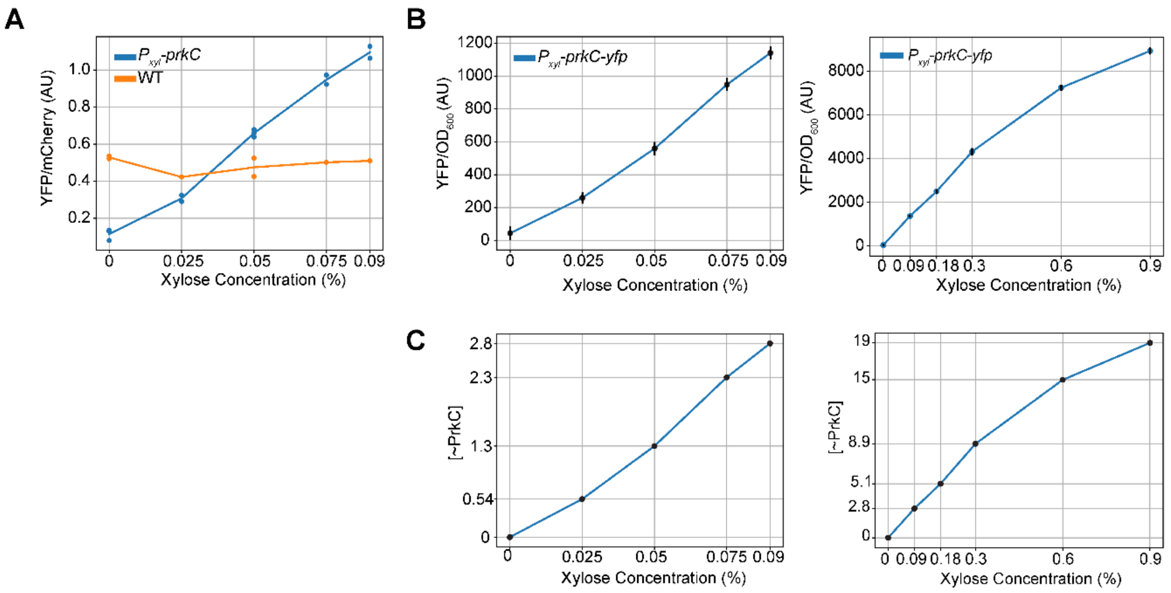


**Figure S3: Complementation and calibration of xylose-inducible PrkC.**

A) **Comparison of inducibly and natively expressed *prkC*.** Kinase activity (YFP/mCherry) of inducible $P_{xylA}$-*prkC* (blue) and natively expressed *prkC* (orange) as a function of xylose concentration reported by LacI~P. Complementing expression of inducible *prkC* ($[PrkC_{inducible}]$ ~ 1) occurs at ~ 0.04% xylose.

B) **Transcriptional reporter for xylose-inducible *prkC*.** Normalized fluorescence ($YFP/OD_{600}$) of a *yfp* transcriptional reporter $P_{xylA}$-*prkC-yfp* (blue) as a function of xylose concentration. Induction was performed up to 0.09% (left) and 0.9% (right) xylose. This curve is used to determine units of PrkC at each inducer level by normalizing signal at each xylose concentration to signal at 0.04% xylose (C, Materials and Methods). Each data point represents the mean of at least 3 biological replicates ±SEM.

C) **Mapping of inducible PrkC expression into native expression units.** Approximate PrkC expression (~[PrkC]) as a function of xylose concentration up to 0.09% (left) and 0.9% (right), where approximately equivalent to native levels are at ~1.

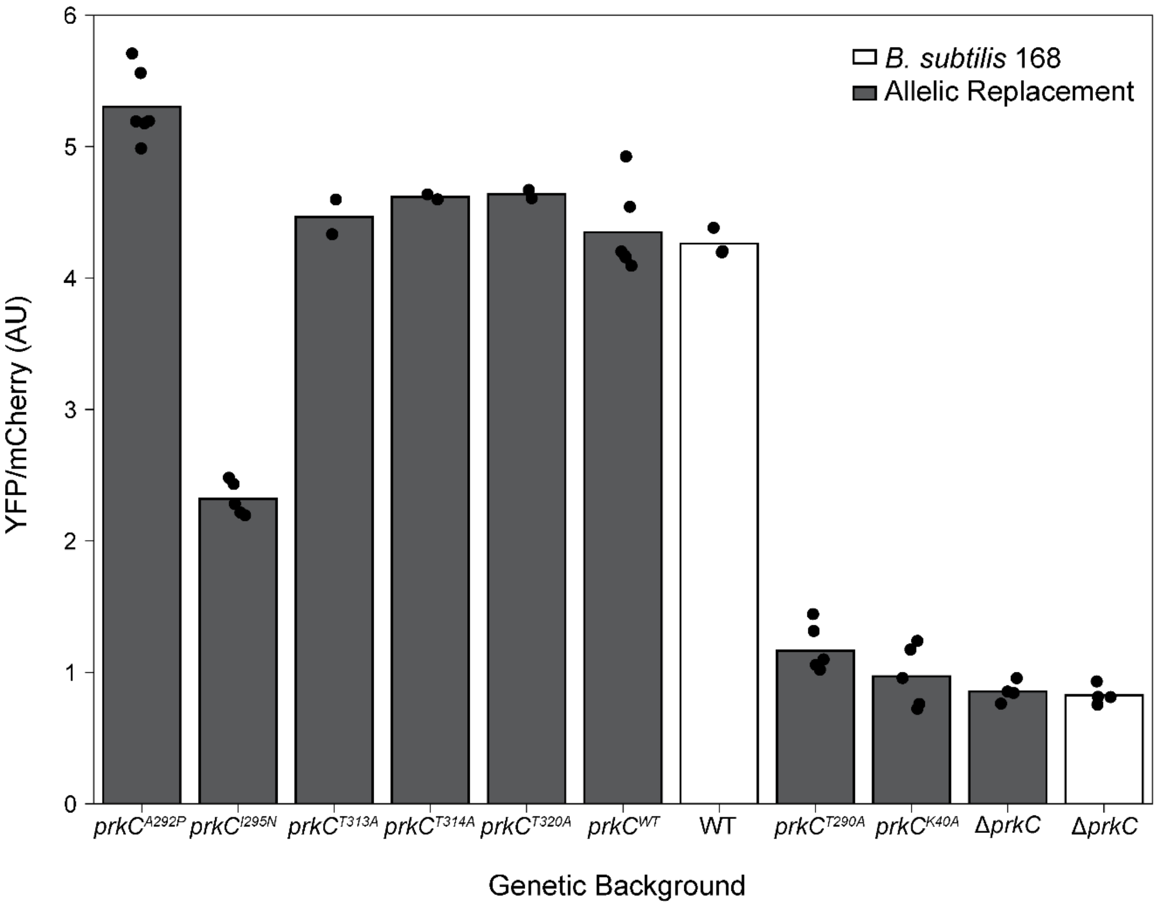


**Figure S4: Kinase activity of allelic replacement strains.** Kinase activity (YFP/mCherry) in log phase of allelic replacement *prkC* variants (grey) compared to WT (white) reported by LacI~P in an otherwise Δ*prpC* background.

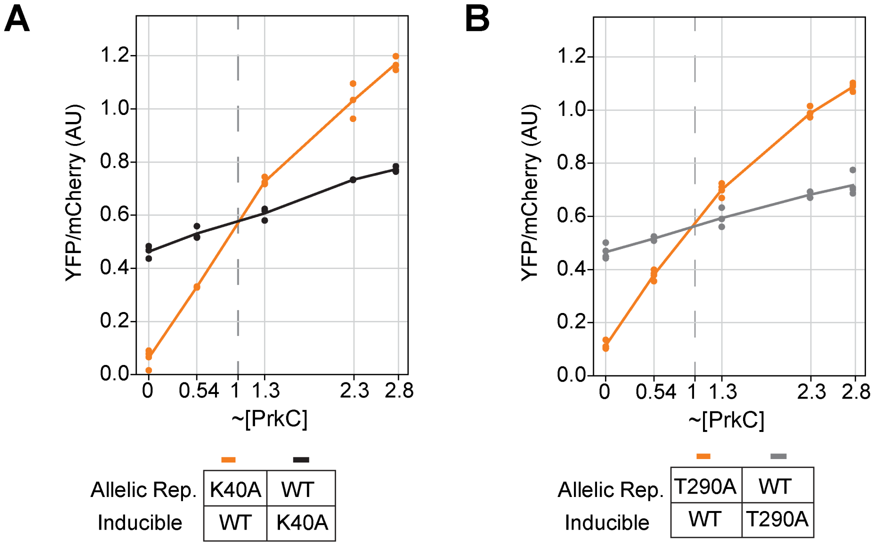


**Figure S5: Total kinase activity is independent of the genetic locus used for *prkC* expression.** Detailed comparison of data in Fig. 4. Expression of *prkC*$^{WT}$ with *prkC*$^{K40A}$ (A) or *prkC*$^{T290A}$ (B) in both possible configurations: *prkC*$^{WT}$ from the inducible promoter and the variant from the native promoter (orange), or with the variant under inducible expression and *prkC*$^{WT}$ from the native promoter (black/grey). Kinase activity (YFP/mCherry) is plotted as a function of inducible PrkC expression level (~[PrkC]). Approximately equal kinase activity occurs in either configuration, where the inducible kinase is expressed at ~[PrkC] = 1 (vertical dashed line).

### Quantitative model of kinase activation

We developed a simple mathematical model to help interpret our quantitative experimental results in the context of a model for kinase signaling pathway activation. The main contributions of this model include

- **Mechanistic validation of feedback**: Demonstrates that autoactivation feedback is the most likely cause for nonlinear activation of kinase variants.
- **Quantitative activation thresholds**: Estimates the expression level at which each variant transitions from low- to high-activity states, explaining feedback-driven behavior.
- **Parameter separation**: Separates catalytic efficiency from activation potential, allowing individual parameter estimation for the substrate and trans-activating kinases.
- **Explaining PrkC$^{\mathbf{K40A}}$ behavior**: Shows that the low activity of PrkC$^{\mathrm{K40A}}$ arises from feedback suppression, with parameter estimation suggesting that a 3-fold decrease in activation potential relative to PrkC$^{\mathrm{WT}}$ is sufficient to suppress its activity at up to nearly 3-fold physiological levels of expression.
- **Global mechanistic consistency**: Confirms that all experimental data, from both single-variant and co-expression experiments, and in the presence of the phosphatase PrpC, are quantitatively consistent with the proposed feedback-regulated mechanism of PrkC activation.

#### Fluorescent reporter of kinase activity

In our experiments, the LacIP kinase activity reporter is used to report kinase activity via fluorescence output. LacIP is believed to exist in two states: unphosphorylated (U) and phosphorylated (P). In the U state, LacIP binds DNA upstream of a fluorescent reporter gene (*yfp*) and represses its transcription, while in the P state it cannot bind DNA and the repression is relieved. *mCherry* is expressed constitutively and is used for normalization (see Materials & Methods). We assume that under the experimental conditions the fluorescence signal $F$ (YFP/mCherry) is linearly proportional to the level of the phosphorylated (deactivated) repressor,

$$F = \gamma L_P \tag{1}$$

where $L_P$ is the steady-state concentration of phosphorylated LacIP and $\gamma$ maps the rate of transcription from the *yfp* promoter to measured fluorescence.

With a fixed total concentration $L_T$ of LacIP, the concentration $L_P$ of high activity reporters can be determined from the balance between the phosphorylation/de-phosphorylation reactions

$$L_P = v_L L_U = v_L(L_T - L_P) \tag{2}$$

yielding

$$F = \sigma \frac{v_L}{1 + v_L} \simeq \sigma v_L \tag{3}$$

where $\sigma = \gamma L_T$. In the last step, we assume a strong bias toward phosphorylation in our experiments, which are done in mutants that do not express the phosphatase.

We write the rate of LacIP phosphorylation, $v_L$ as the product of two factors: the phosphorylation susceptibility of LacIP ($\theta_L$) and the total catalytic activity of PrkC kinases in the cell ($v_{\mathrm{Tot}}$):

$$F = \sigma \theta_L v_{\mathrm{Tot}} \; . \tag{4}$$

In our experimental setup, $\theta_L$ is kept constant, as its value is intrinsically determined. In contrast, total activity $v_{\mathrm{Tot}}$ is modulated by mixing different PrkC variants and by controlling their expression levels through an inducible promoter system. Consequently, in all experiments, the measured steady-state fluorescence levels are proportional to $v_{\mathrm{Tot}}$ with a fixed proportionality constant, which will be omitted from the rest of the discussion.

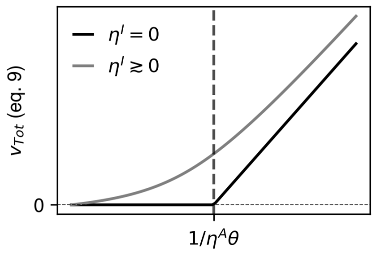


Figure T1: Expected threshold-like form of fluorescence (F) as function of total kinase concentration. The non-linear thresholding response is due to autoactivation.

#### Autoactivation of the kinase

Many of the strains studied in this work express a single PrkC variant. This can be either the wild-type kinase or one of its variants. PrkC can be phosphorylated at multiple sites, including on its activation loop and the juxtamembrane IDR. We hypothesize that it can therefore exhibit multiple phosphorylation states. In our model we represent these states by two "macro"-states: a low activity state (L), characterized by a lower catalytic activity $\eta_L$, and a high activity state (H), characterized by the higher activity $\eta_H$. With $x_L$ and $x_H$ as the concentration of PrkC in these two states, respectively, the total kinase activity is given by

$$v_{\text{Tot}} = \eta_H x_H + \eta_L x_L \ . \tag{5}$$

In our experiments we control the total kinase concentration $x_T = x_L + x_H$. We let $x_T = 1$ for expression from the native *prkC* promoter and define other expression levels in these units (see Materials & Methods and Supp. Fig. S3).

Assuming that transitions from the low activity to the high activity state require phosphorylation event(s), the steady-state balance between $x_H$ and $x_L$ is given, similar to (2), by

$$x_H = v(x_T - x_H) \ . \tag{6}$$

As for LacIP, the rate of phosphorylation $v$ is a product of the total kinase activity $v_{\text{Tot}}$ and the collective susceptibility of the relevant phosphorylation sites, which we denote by $\theta$. We therefore insert

$$v = \theta v_{\text{Tot}} = \theta \left[\eta_L x_L + \eta_H x_H\right] \tag{7}$$

in (6) to get

$$x_H = \theta \left[\eta_L (x_T - x_H) + \eta_H x_H\right] (x_T - x_H) \ . \tag{8}$$

The non-linear (quadratic) nature of this equation reflects the positive feedback due to autoactivation.

Finally, we obtain the total kinase activity by solving this equation for $x_H$ and substituting the biologically relevant root (between 0 and $x_T$) into (5)

$$v_{\text{Tot}} = \frac{1}{2}\left[(x_T \eta_H \theta - 1) + \sqrt{(x_T \eta_H \theta - 1)^2 + 4 x_T \eta_L \theta}\right] \ . \tag{9}$$

This expression captures how total kinase activity (measured by reporter fluorescence, (4)) depends on the underlying properties of the kinase, namely its susceptibility to activation ($\theta$) and its activity in the low activity and high activity states ($\eta_L$ and $\eta_H$).

##### Threshold-like behavior of kinase activation

We attribute the unique features of some of the variants in our study, including the observed nonlinearity of PrkC$^{\text{T290A}}$ activity and the undetectable activity of PrkC$^{\text{K40A}}$, to a threshold-like behavior due to weak

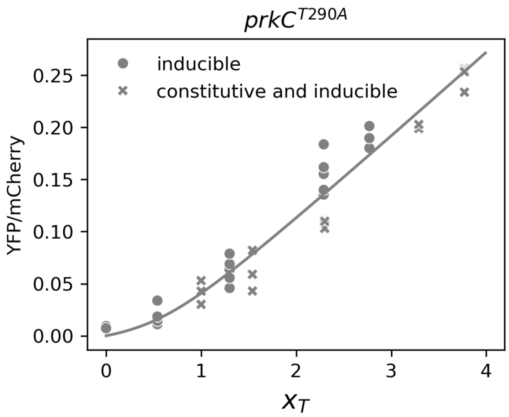


Figure T2: Model fit for PrkC$^{\mathrm{T290A}}$ activity. Kinase activity reporter fluorescence is plotted against total kinase concentration ($x_T$). Symbols are experimental data from fluorescent measurements, and lines are the best fit of the autoactivation model.

autoactivation. To illustrate this behavior, consider the case $\eta_L \ll \eta_H$. In this limit we expand the expression for $v_{\mathrm{Tot}}$ given in (9) to first order:

$$v_{\mathrm{Tot}} \approx [x_T \eta_H \theta - 1]_+ + \frac{x_T \eta_L \theta}{|x_T \eta_H \theta - 1|} \tag{10}$$

where $[x]_+ \equiv \max(0, x)$ is the ramp function (ReLU in machine learning). This expression shows the nonlinear threshold-like dependence of kinase activity on its total concentration $x_T$. When $x_T < {}^1/_{\theta\eta_H}$ the leading term vanishes, and $v$ grows slowly with $x_T$ (or remains zero in the extreme case $\eta_L$=0). Conversely, when $x_T > {}^1/_{\theta\eta_H}$ the leading term dominates, and $v_L$ grows linearly with the steep slope $\eta_H$ (Fig. T1).

#### Co-expression of two kinase variants

Our results show that autoactivation of PrkC occurs in trans. When more than one variant of *prkC* is expressed, all variants contribute to the total kinase activity $v_{\mathrm{Tot}}$. In our experiments, some strains co-express two different kinase variants, and in these cases (5) is replaced by

$$v_{\mathrm{Tot}} = \left[\eta_H^1 x_H^1 + \eta_L^1 (x_T^1 - x_H^1)\right] + \left[\eta_H^2 x_H^2 + \eta_L^2 (x_T^2 - x_H^2)\right] \tag{11}$$

with the superscript indicating the kinase variant. Each variant maintains its own steady-state balance between low activity and high activity states:

$$x_H^1 = v_1 (x_T^1 - x_H^1) \ , \qquad x_H^2 = v_2 (x_T^2 - x_H^2) \tag{12}$$

with

$$v_1 = \theta_1 v_{\mathrm{Tot}} \ , \qquad v_2 = \theta_2 v_{\mathrm{Tot}} \ . \tag{13}$$

Plugging (11) and (13) into (12) results in a pair of coupled quadratic equations. Although an analytical solution is possible, identifying and interpreting the biologically relevant root is neither straightforward nor particularly insightful. Therefore, we opt for a numerical solution, which offers both simplicity and accuracy.

#### Parameter estimation

To evaluate the applicability of our simple model to PrkC function in vivo and to explore underlying mechanisms, we inferred model parameters using the data presented in Fig. 4A of the main text and used it to interpret data in Fig. 4C. It is important to note that our model is a highly simplified "toy" model, and

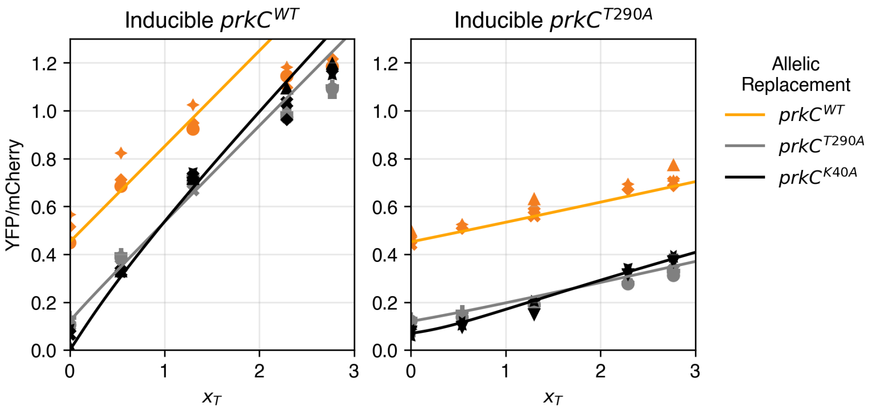


Figure T3: Comparison of model prediction with experimental datasets in co-expression experiments. Symbols are experimental data as in Fig. 4C of the main text, lines are model predictions with the parameters detailed in the Table T1. Colors indicate the expressed allelic replacement variant.

the parameter values are only meant to compare different mechanisms. All parameters should be considered "dimensionless" except $\sigma$, which carries arbitrary fluorescence units.

We first considered only cells that express a single variant. In these cases we can only infer the combination $\theta\eta_H$ and $\theta\eta_L$ and cannot infer the value of these parameters separately. For cells that only express *prkC*$^{T290A}$, which shows strong nonlinear behavior as a function of total kinase concentration $x_T$, we find $\theta_{\mathrm{T290A}}\eta_H \simeq 0.9(\pm 0.1)$ and $\theta_{\mathrm{T290A}}\eta_L = 0.3(\pm 0.1)$ with $\sigma = 0.1(\pm 0.02)$ (Fig. T2).

For PrkC$^{\mathrm{WT}}$, fluorescence increases almost linearly with $x_T$. This allows us to infer $\theta_{\mathrm{WT}}\eta_H = 3.1(\pm 0.3)$, and estimate that $\theta_{\mathrm{WT}}\eta_L > 2$. These results suggest that IDR phosphorylation enhances activity in both the low activity and high activity states. For example, it is possible that an unphosphorylated IDR physically interferes with substrate binding in both states.

PrkC$^{\mathrm{K40A}}$ remains inactive in the range of $x_T$ explored in our experiment. Within our model, this implies $\eta_L = 0$ and suggests that $x_T$ remains below the threshold at all expression levels, implying $\theta_{\mathrm{K40A}}\eta_H < 1/3$. If the K40A mutation inhibits the transition to the high activity state but not the activity of the kinase in that state, then $\eta_H$ would assume the same value as for the wild-type, and $\theta_{\mathrm{K40A}} \leq 0.11$.

To validate the model and the estimated parameter values, we compared the model predictions with the experimental data shown in Fig. 4C of the main text, which considers cells that co-express two kinase variants. Here we need to assign values to all $\eta$ and $\theta$ separately. We have the freedom to set the value of one of the $\theta$'s to any arbitrary number, which will simply set the scale for all the other parameters, so we set $\theta_{\mathrm{WT}} = 1$. The argument above leads us to set $\theta_{\mathrm{K40A}} = 0.11$. With these values, we find that letting $\theta_{\mathrm{T290A}} = 1$ yields a satisfying fit to the data. Fig. T3 compares the model predictions and the experimental data for cells that express *prkC*$^{WT}$ or *prkC*$^{T290A}$ from the inducible promoter with other variants co-expressed from the native promoter via allelic replacement. The quality of this fit provides strong support of our model. In particular, it captures well the trans-activation effect, which serves to activate PrkC$^{\mathrm{K40A}}$ in the presence of other variants.

As described in the main text, we observe a shift in the activity of some variants at very high levels of PrkC induction. This shift could be due to saturation of the reporter substrate, dimerization or multimerization of the kinase, or other physical constraints, and is not captured by the model.

#### Exploring mechanisms of activation

One interesting feature, observed both experimentally and in model simulations, occurs in cells that co-express *prkC*$^{T290A}$ and *prkC*$^{K40A}$. When we compare cells that express *prkC*$^{T290A}$ from the native promoter and *prkC*$^{K40A}$ or a second copy of *prkC*$^{T290A}$ from the inducible promoter, we find that the activity of the

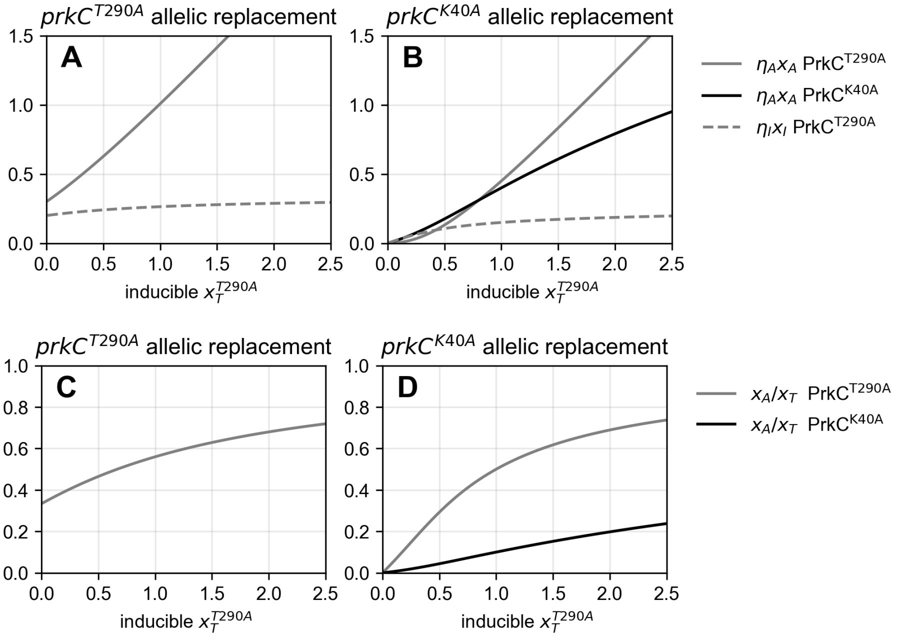


Figure T4: Individual contributions of kinase variants to total activity. Kinase activities of different variants, in cells expressing *prkC*$^{T290A}$ (solid lines) or *prkC*$^{K40A}$ (dashed lines) from the native promoter.

former is lower at low levels of induction but higher at high levels (black and gray lines in right panel of Fig.T3).

To explain this, we look at the contributions of the two variants to the total activity (Fig. T4AB). By itself, PrkC$^{\mathrm{K40A}}$ kinases cannot autoactivate, which is why cells that carry constitutive *prkC*$^{K40A}$ show no activity at no inducer, while cells that carry *prkC*$^{T290A}$ show baseline activity. As the concentration of PrkC$^{\mathrm{T290A}}$ grows, in cells that only carry *prkC*$^{T290A}$ this population autoactivates itself, and the fraction of high activity kinases in the population grows (Fig. T4C), creating the nonlinear increase in activation (Fig. T4A). In cells expressing *prkC*$^{K40A}$ from the native promoter, a growing PrkC$^{\mathrm{T290A}}$ population also activates PrkC$^{\mathrm{K40A}}$, which has higher activity once high activity ($\eta_H^{\mathrm{K40A}} > \eta_H^{\mathrm{T290A}}$), resulting in overall higher activity. Moreover, the highly active PrkC$^{\mathrm{K40A}}$ activates the PrkC$^{\mathrm{T290A}}$ population (compare gray lines in Fig. T4C and T4D), adding another contribution to the overall activity (Fig. T4B).

#### Quantitative effects of the phosphatase

As seen above, the activity of the the wildtype kinase in the absence of the phosphatase is nearly linear in its total concentration, and does not display the threshold effect. In contrast, titration of *prkC*$^{WT}$, as well as the hyper-active *prkC*$^{A292P}$, in the presence of the phosphatase shows a highly nonlinear dependence on their total concentrations. We hypothesized that this response reflects the same underlying autoactivation model, where the phosphatase effectively reduces the phosphorylation rates $\eta_L$ and $\eta_H$.

Indeed, fitting the data from Fig. 5B of the main text to our model's prediction (9) provides strong support to this hypothesis (Fig. T5). For *prkC*$^{WT}$ we find $\theta\eta_L = 0.006$ and $\theta\eta_H = 0.12$, while for *prkC*$^{A292P}$ we have $\theta\eta_L = 0.009$ and $\theta\eta_H = 0.20$.

| variant | $prpC$ | parameter | value | comments |
|---|---|---|---|---|
| | | $\eta^L$ | $> 2$ | |
| PrkC$^{\mathrm{WT}}$ | – | $\eta^H$ | 3.1 | |
| | | $\theta$ | $\simeq 1$ | |
| PrkC$^{\mathrm{WT}}$ | + | $\theta\eta^L$ | 0.006 | |
| | | $\theta\eta^H$ | 0.12 | |
| | | $\eta^L$ | 0.3 | |
| PrkC$^{\mathrm{T290A}}$ | – | $\eta^H$ | 0.9 | |
| | | $\theta$ | 1 | by definition |
| | | $\eta^L$ | 0 | |
| PrkC$^{\mathrm{K40A}}$ | – | $\eta^H$ | $< 3$ | |
| | | $\theta$ | $\simeq 0.1$ | |
| PrkC$^{\mathrm{A292P}}$ | + | $\theta\eta^L$ | 0.009 | |
| | | $\theta\eta^H$ | 0.20 | |

Table T1: Parameter values

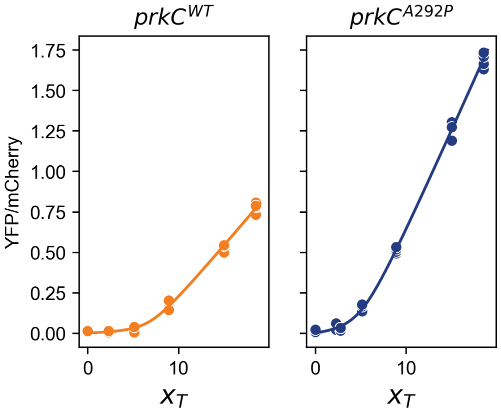


Figure T5: Model fit for activity in the presence of the phosphatase. Reporter fluorescence is plotted against total kinase concentration ($x_T$). Symbols are experimental data from fluorescence measurements (Fig. 5B of the main text), and lines are the best fit to the autoactivation model.

#### Materials and Methods

##### IDR conservation analysis

Sequences of bacterial and archaeal kinases whose catalytic domains were previously aligned[1], along with additional Hanks-type Ser/Thr kinases from bacterial species selected based on the presence of a disordered domain[2,3], were retrieved from UniProtKB[4] via API and aligned using MAFFT based on the full kinase sequences. The alignment region corresponding to the IDR of *B. subtilis* PrkC was then extracted. Conservation at each residue position was quantified as the fraction of sequences containing the most commonly appearing residue at that position. An alignment plot was generated using Matplotlib in Python. The presence of a juxtamembrane domain was predicted using DeepTMHMM[5] and Alphafold pLDDT scores[2,3].

A protein sequence similarity search for *B. subtilis* PrkC was performed using BLASTP[6] against the NCBI non-redundant (nr) protein database using a default E-value threshold of 1e-5. The top 500 hits were analyzed for sequence identity and functional annotations using the InterProScan 5 API[7]. Amino acid sequence variation was assessed by hierarchical clustering of sequences based on pairwise string distance. Sequence logos were generated using Logomaker from position probability matrices.

##### Plasmid construction

Lists of oligos and plasmids used in this work can be found in Supplementary Data (Tables S3, S4). Oligos for plasmid construction were synthesized by Genewiz. PCR was performed using Q5 High-Fidelity 2X Master Mix (NEB #M0492L) and purified using the Monarch PCR & DNA Cleanup Kit (NEB #T1030, #T1130) or Monarch DNA Gel Extraction Kit (NEB #T1020, #T1120). Golden Gate cloning was done with PaqCI enzyme (NEB #R0745) or BsaI enzyme (NEB #E1602). Plasmids were cloned into NEB 5-alpha competent *E. coli* (NEB C2987H) or Turbo competent *E. coli* (NEB C2984H) (see Supplementary Table S2), with selection and propagation on LB lennox ampicillin 100 µg/ml (50 µg/ml for pEL53). Plasmid DNA was purified using the Monarch Plasmid DNA Miniprep Kit (NEB #T1010, #T1110). Construction was verified by Sanger sequencing or whole plasmid sequencing by Genewiz. See Table S3 for details of plasmid construction.

**Strain construction**

Lists of strains used in figures and in this study can be found in Supplementary Data (Table S1, S2). *B. subtilis* strains were constructed by transformation of plasmid DNA, genomic DNA, or PCR products into *B. subtilis* made competent using the 2-step method[8] and selection on LB lennox agar plates containing appropriate antibiotic (5 µg/mL chloramphenicol, 10 µg/mL kanamycin, 10 µg/mL tetracycline, or MLS). Integration at the *sacA* locus was verified by lack of growth on sucrose agar plates[8] and integration at the *ganA and prkC* loci were verified by PCR. All strains constructed by allelic replacement at the *prkC* and *prpC* loci were verified by Sanger sequencing PCR amplicons of the mutated locus region from genomic DNA. Whole-genome sequencing was also performed by Plasmidsaurus on genomic DNA of a selection of allelic replacement strains (NGB379, NGB380, NGB427, NGB428). Genomic DNA was purified using the Wizard Genomic DNA Purification Kit (Promega A1120).

**Growth media and culture conditions**

*B. subtilis* strains were streaked for single colonies from frozen stocks on LB lennox agar plates and grown at 37°C (16-18 hours) for liquid experiments or for observation of lysis on plates (18-24 hours). Agar plates were imaged with an Epson Perfection V600 scanner (Epson B11B198011).

Liquid cultures were inoculated from single colonies in 2 mL S7-glucose growth media, LB lennox broth, or CH media[8]. S7-glucose growth media was prepared by supplementing the

MOPS Minimal Media kit (Teknova M2106) with 0.1% w/v L-glutamic acid (Sigma-Aldrich 49621) and 40 μg/mL L-tryptophan (Sigma-Aldrich T0254) as described previously[9,10]. For antibiotic sensitivity experiments, single colonies were grown in 2 mL S7-glucose media shaking at 37°C to mid-log phase. MIC test-strip experiments were performed by spreading dense culture on S7-agar plates, placing cefotaxime MIC test-strips (Liofilchem 92006) on the plate, and reading MIC values after 18-20 hours as previously described[9]. Measurements of gene expression in liquid culture were performed in 96-well microplates. Cultures were inoculated from single colonies and grown to mid-log in 2 mL of growth media with aeration at 37°C. They were diluted 1:30 in fresh media, supplemented with xylose as indicated (0, 0.025, 0.05, 0.075, and 0.09% final concentrations), to a final volume of 150 μL per well in a microplate in triplicate. Microplates were incubated at 37°C with continuous shaking in a BioTek Cytation 5 plate reader for $OD_{600}$ and luminescence reads or in a BioTek Synergy Neo2 plate reader for $OD_{600}$, YFP fluorescence (500/27 ex, 540/25 em), and mCherry fluorescence (560/20 ex, 620/15 em) reads.

**Fluorescence and luminescence data processing**

Background subtraction and normalization: For each timepoint measurement in 96-well microplate experiments, sample $OD_{600}$ and reporter signals (luminescence, YFP, or mCherry) were corrected by subtracting values from media-only wells. Corrected reporter signals were then normalized by corresponding corrected $OD_{600}$.

Autofluorescence correction: Normalized reporter signals were further corrected by subtracting values measured for Bs168 control samples in each signal channel. For fluorescence experiments, YFP signals were additionally normalized by constitutive mCherry signals.

Triplicate wells were averaged at each measurement timepoint. Final values for analysis were calculated within defined OD600 range (~0.8-0.9 for luminescence, ~0.35-0.45 for fluorescence).

For experiments using a xylose-inducible system, YFP/mCherry values were calculated as described above for each xylose concentration. Autofluorescence correction was applied separately for each xylose concentration using the corresponding Bs168 control values.

For experiments conducted with both xylose and cefotaxime, cultures were first grown in CH media with xylose in a 96-well microplate in a plate reader to early-log phase ($OD_{600}$ ~0.3). Cefotaxime was then added at a 1:100 dilution to a final concentration of 0.05 µg/mL and cultures were further incubated at 37°C with continuous shaking for 3 hours. $OD_{600}$ and fluorescence signals were measured and YFP/mCherry values were calculated as described above at $OD_{600}$ range ~0.5-0.6.

**Quantification of inducible *prkC* expression**

Complementing expression of inducible *prkC* was empirically determined as the xylose concentration at which the inducible expression level from *ganA::*$P_{xyl}$-*prkC* (NGB249) matched expression levels from the native promoter (NGB57) in an otherwise Δ*prkC* background, as measured by YFP/mCherry LacI~P signal (~0.04% xylose, Supplementary Figure S3A). This concentration was denoted as 1 unit of PrkC.

To determine relative expression levels from the $P_{xyl}$ promoter, fluorescence of a $P_{xyl}$-*yfp* transcriptional reporter (NGB539) was measured at each xylose concentration and processed as described above (Supplementary Figure S3B). Signals were background-subtracted using the 0% xylose measurement. Relative [$PrkC_{inducible}$] was calculated by dividing each value by the signal at the complementing xylose concentration (Supplementary Figure S3C).

**S1 Table: Strains used in data figures**

| Figure | Panel | Strain |
|---|---|---|
| 2 | A | Bs168, JDB1773, JDB1775, NGB493, NGB494, NGB495, NGB496, NGB497, NGB527, NGB528 |
| | B | NGB367, NGB369, NGB372, NGB379, NGB380, NGB382 |
| | C | NGB367, NGB369, NGB372, NGB379, NGB380, NGB382, NGB427, NGB428, NGB431, NGB432 |
| 3 | B | NGB498, NGB499, NGB500, NGB547 |
| | C | NGB379, NGB380, NGB382, NGB385 |
| 4 | A | NGB249, NGB261 |
| | B | NGB249, NGB261, NGB265, NGB535, NGB410, NGB526, NGB408, NGB530 |
| | D | NGB436, NGB441, NGB454, NGB435, NGB438, NGB456, NGB437, NGB440, NGB455, NGB560, NGB561, NGB562 |
| 5 | A | NGB437, NGB441, NGB561 |
| | B | NGB249, NGB261, NGB408<br>NGB248, NGB260, NGB407 |
| S2 | B | NGB493, NGB494, NGB495, NGB527 |
| S3 | A | NGB57, NGB249 |
| | B, C | NGB539 |
| S4 | | NGB57, NGB58, NGB379, NGB380, NGB382, NGB385, NGB420, NGB421, NGB431, NGB432, NGB467 |
| S5 | A | NGB435, NGB454 |
| | B | NGB436, NGB440 |

**S2 Table: Strains used in this study**

| Strain | Genotype | Construction | Source |
|---|---|---|---|
| ***E. coli*** | | | |
| 5-alpha | *fhuA2Δ(argF-lacZ)U169 phoA glnV44 Φ80Δ(lacZ)M15 gyrA96 recA1 relA1 endA1 thi-1 hsdR17* | | New England Biolabs |
| Turbo | F' *proA*$^{+}$*B*$^{+}$ *lacI*$^{q}$ *ΔlacZM15 / fhuA2 Δ(lac-proAB) glnV galK16 galE15 R(zgb-210::Tn10)*Tet$^{S}$ *endA1 thi-1 Δ(hsdS-mcrB)5* | | New England Biolabs |
| ***B. subtilis*** | | | |
| Bs168 | 168 *trpC2* (WT) | | Lab stock & [11] |
| ELB203 | *sacA::*$P_{pdaC}$*-luxABCDE cm* | | Lab stock & [11] |
| ELB204 | *sacA::*$P_{iseA}$*-luxABCDE cm* | | Lab stock & [11] |
| JDB1773 | 168 *trpC2* Δ*prpC* | | Lab stock & [11] |
| JDB1774 | 168 *trpC2* Δ*prkC* | | Lab stock & [11] |
| JDB1775 | 168 *trpC2* Δ*(prpC-prkC)* | | Lab stock & [11] |
| JDB2482 | Δ*prpC::tet* | | Gift from Jonathan Dworkin |
| NGB52 | *sacA::(*$P_{veg}$*-mCherry,* $P_{P1}$*-lacIP,* $P_{lac}$*-yfp) cm* | Integration of pNG21 into Bs168 | This study |
| NGB57 | Δ*prpC sacA::(*$P_{veg}$*-mCherry,* $P_{P1}$*-lacIP,* $P_{lac}$*-yfp) cm* | Transformation of NGB52 into JDB1773 | This study |
| NGB58 | Δ*(prpC-prkC) sacA::(*$P_{veg}$*-mCherry,* $P_{P1}$*-lacIP,* $P_{lac}$*-yfp) cm* | Transformation of NGB52 into JDB1775 | This study |
| NGB244 | *ganA::*$P_{xyl}$*-prkC mls* | Integration of pNG130 into Bs168 | This study |
| NGB239 | Δ*prkC sacA::(*$P_{veg}$*-mCherry,* $P_{P1}$*-lacIP,* $P_{lac}$*-yfp) cm* | Transformation of NGB52 into JDB1774 | This study |
| NGB245 | *ganA::*$P_{xyl}$*-prkC*$^{T290A}$ *mls* | Integration of pNG131 into Bs168 | This study |
| NGB246 | *ganA::*$P_{xyl}$*-prkC*$^{K40A}$ *mls* | Integration of pNG135 into Bs168 | This study |
| NGB248 | Δ*prkC sacA::(*$P_{veg}$*-mCherry,* $P_{P1}$*-lacIP,* $P_{lac}$*-yfp) cm, ganA::*$P_{xyl}$*-prkC mls* | Transformation of NGB244 into NGB239 | This study |
| NGB249 | Δ*(prpC-prkC) sacA::(*$P_{veg}$*-mCherry,* $P_{P1}$*-lacIP,* $P_{lac}$*-yfp) cm, ganA::*$P_{xyl}$*-prkC mls* | Transformation of NGB244 into NGB58 | This study |
| NGB260 | Δ*prkC sacA::(*$P_{veg}$*-mCherry,* $P_{P1}$*-lacIP,* $P_{lac}$*-yfp) cm, ganA::*$P_{xyl}$*-prkC*$^{T290A}$ *mls* | Transformation of NGB245 into NGB239 | This study |
| NGB261 | Δ*(prpC-prkC) sacA::(*$P_{veg}$*-mCherry,* $P_{P1}$*-lacIP,* $P_{lac}$*-yfp) cm, ganA::*$P_{xyl}$*-prkC*$^{T290A}$ *mls* | Transformation of NGB245 into NGB58 | This study |
| NGB265 | Δ*(prpC-prkC sacA::(*$P_{veg}$*-mCherry,* $P_{P1}$*-lacIP,* $P_{lac}$*-yfp) cm, ganA::*$P_{xyl}$*-prkC*$^{K40A}$ *mls* | Transformation of NGB246 into NGB58 | This study |

| | | | |
|---|---|---|---|
| NGB328 | *prkC::kan* | Integration of pNG146 into Bs168 | This study |
| NGB342 | *prkC::prkC*$^{K40A}$ *kan* | Integration of pNG154 into Bs168 | This study |
| NGB345 | *prkC::prkC*$^{T290A}$ *kan* | Integration of pNG152 into Bs168 | This study |
| NGB367 | *sacA::(*$P_{veg}$*-mCherry,* $P_{P1}$*-lacIP,* $P_{lac}$*-yfp) cm, prkC::kan* | Transformation of NGB328 into NGB52 | This study |
| NGB369 | *sacA::(*$P_{veg}$*-mCherry,* $P_{P1}$*-lacIP,* $P_{lac}$*-yfp) cm, prkC::prkC*$^{K40A}$ *kan* | Transformation of NGB342 into NGB52 | This study |
| NGB372 | *sacA::(Pveg-mCherry, P1-lacIP, Plac-yfp) cm, prkC::prkC*$^{T290A}$ *kan* | Transformation of NGB345 into NGB52 | This study |
| NGB379 | *sacA::(*$P_{veg}$*-mCherry,* $P_{P1}$*-lacIP,* $P_{lac}$*-yfp) cm, prkC::kan, ΔprpC::tet* | Integration of Δ*prpC::tet* PCR product from JDB2482 genomic DNA using primers NGp67 and NGp196 into NGB367 | This study |
| NGB380 | *sacA::(*$P_{veg}$*-mCherry,* $P_{P1}$*-lacIP,* $P_{lac}$*-yfp) cm, prkC::prkC*$^{T290A}$ *kan, ΔprpC::tet* | Integration of Δ*prpC::tet* PCR product from JDB2482 genomic DNA using primers NGp67 and NGp196 into NGB372 | This study |
| NGB382 | *sacA::(*$P_{veg}$*-mCherry,* $P_{P1}$*-lacIP,* $P_{lac}$*-yfp) cm, prkC::prkC*$^{K40A}$ *kan, ΔprpC::tet* | Integration of Δ*prpC::tet* PCR product from JDB2482 genomic DNA using primers NGp67 and NGp196 into NGB369 | This study |
| NGB385 | *sacA::(*$P_{veg}$*-mCherry,* $P_{P1}$*-lacIP,* $P_{lac}$*-yfp) cm, Δ(prpC-prkC)::kan* | Integration of pNG162 into NGB52 | This study |
| NGB395 | *prkC::prkC*$^{A292P}$ *kan* | Integration of pNG168 into Bs168 | This study |
| NGB396 | *prkC::prkC*$^{I295N}$ *kan* | Integration of pNG171 into Bs168 | This study |
| NGB397 | *ganA::*$P_{xyl}$*-prkC*$^{A292P}$ *mls* | Integration of pNG167 into Bs168 | This study |
| NGB401 | *sacA::(*$P_{veg}$*-mCherry,* $P_{P1}$*-lacIP,* $P_{lac}$*-yfp) cm, prkC::prkC*$^{A292P}$ *kan* | Transformation of NGB395 into NGB52 | This study |
| NGB402 | *sacA::(*$P_{veg}$*-mCherry,* $P_{P1}$*-lacIP,* $P_{lac}$*-yfp) cm, prkC::prkC*$^{I295N}$ *kan* | Transformation of NGB396 into NGB52 | This study |
| NGB407 | *ΔprkC sacA::(*$P_{veg}$*-mCherry,* $P_{P1}$*-lacIP,* $P_{lac}$*-yfp) cm, ganA::*$P_{xyl}$*-prkC*$^{A292P}$ *mls* | Transformation of NGB397 into NGB239 | This study |
| NGB408 | *Δ(prpC-prkC) sacA::(*$P_{veg}$*-mCherry,* $P_{P1}$*-lacIP,* $P_{lac}$*-yfp) cm, ganA::*$P_{xyl}$*-prkC*$^{A292P}$ *mls* | Transformation of NGB397 into NGB58 | This study |
| NGB416 | *prkC::prkC*$^{T313A}$ *kan* | Integration of pNG172 into Bs168 | This study |

| | | | |
|---|---|---|---|
| NGB417 | *prkC::prkC$^{T320A}$ kan* | Integration of pNG173 into Bs168 | This study |
| NGB420 | *sacA::(P$_{veg}$-mCherry, P$_{P1}$-lacIP, P$_{lac}$-yfp) cm, prkC::prkC$^{A292P}$ kan, ΔprpC::tet* | Integration of Δ*prpC::tet* PCR product from JDB2482 genomic DNA using primers NGp67 and NGp196 into NGB401 | This study |
| NGB421 | *sacA::(P$_{veg}$-mCherry, P$_{P1}$-lacIP, P$_{lac}$-yfp) cm, prkC::prkC$^{I295N}$ kan, ΔprpC::tet* | Integration of Δ*prpC::tet* PCR product from JDB2482 genomic DNA using primers NGp67 and NGp196 into NGB402 | This study |
| NGB427 | *sacA::(P$_{veg}$-mCherry, P$_{P1}$-lacIP, P$_{lac}$-yfp) cm, prkC::prkC$^{T313A}$ kan* | Transformation of NGB416 into NGB52 | This study |
| NGB428 | *sacA::(Pveg-mCherry, P1-lacIP, Plac-yfp) cm, prkC::prkC$^{T320A}$ kan* | Transformation of NGB417 into NGB52 | This study |
| NGB431 | *sacA::(P$_{veg}$-mCherry, P$_{P1}$-lacIP, P$_{lac}$-yfp) cm, prkC::prkC$^{T313A}$ kan, ΔprpC::tet* | Integration of Δ*prpC::tet* PCR product from JDB2482 genomic DNA using primers NGp67 and NGp196 into NGB427 | This study |
| NGB432 | *sacA::(P$_{veg}$-mCherry, P$_{P1}$-lacIP, P$_{lac}$-yfp) cm, prkC::prkC$^{T320A}$ kan, ΔprpC::tet* | Integration of Δ*prpC::tet* PCR product from JDB2482 genomic DNA using primers NGp67 and NGp196 into NGB428 | This study |
| NGB435 | *sacA::(P$_{veg}$-mCherry, P$_{P1}$-lacIP, P$_{lac}$-yfp) cm, prkC::kan, ΔprpC::tet, ganA::P$_{xyl}$-prkC$^{K40A}$ mls* | Integration of pNG135 into NGB379 | This study |
| NGB436 | *sacA::(P$_{veg}$-mCherry, P$_{P1}$-lacIP, P$_{lac}$-yfp) cm, prkC::prkC$^{T290A}$ kan, ΔprpC::tet, ganA::P$_{xyl}$-prkC mls* | Integration of pNG130 into NGB380 | This study |
| NGB437 | *sacA::(P$_{veg}$-mCherry, P$_{P1}$-lacIP, P$_{lac}$-yfp) cm, prkC::prkC$^{T290A}$ kan, ΔprpC::tet, ganA::P$_{xyl}$-prkC$^{T290A}$ mls* | Integration of pNG131 into NGB380 | This study |
| NGB438 | *sacA::(P$_{veg}$-mCherry, P$_{P1}$-lacIP, P$_{lac}$-yfp) cm, prkC::prkC$^{T290A}$ kan, ΔprpC::tet, ganA::P$_{xyl}$-prkC$^{K40A}$ mls* | Integration of pNG135 into NGB380 | This study |
| NGB440 | *sacA::(P$_{veg}$-mCherry, P$_{P1}$-lacIP, P$_{lac}$-yfp) cm, prkC::kan, ΔprpC::tet, ganA::P$_{xyl}$-prkC$^{T290A}$ mls* | Integration of pNG131 into NGB379 | This study |
| NGB441 | *sacA::(P$_{veg}$-mCherry, P$_{P1}$-lacIP, P$_{lac}$-yfp) cm, prkC::kan, ΔprpC::tet, ganA::P$_{xyl}$-prkC mls* | Integration of pNG130 into NGB379 | This study |
| NGB445 | *prkC::prkC$^{T314A}$ kan* | Integration of pNG177 into Bs168 | This study |

| | | | |
|---|---|---|---|
| NGB454 | *sacA::($P_{veg}$-mCherry, $P_{P1}$-lacIP, $P_{lac}$-yfp) cm, prkC::prkC$^{K40A}$ kan, ΔprpC::tet, ganA::$P_{xyl}$-prkC mls* | Integration of pNG130 into NGB382 | This study |
| NGB455 | *sacA::(Pveg-mCherry, P1-lacIP, Plac-yfp) cm, prkC::prkC$^{K40A}$ kan, ΔprpC::tet, ganA::$P_{xyl}$-prkC$^{T290A}$ mls* | Integration of pNG131 into NGB382 | This study |
| NGB456 | *sacA::($P_{veg}$-mCherry, $P_{P1}$-lacIP, $P_{lac}$-yfp) cm, prkC::prkC$^{K40A}$ kan, ΔprpC::tet, ganA::$P_{xyl}$-prkC$^{K40A}$ mls* | Integration of pNG135 into NGB382 | This study |
| NGB458 | *sacA::($P_{veg}$-mCherry, $P_{P1}$-lacIP, $P_{lac}$-yfp) cm, prkC::prkC$^{T314A}$ kan* | Transformation of NGB445 into NGB52 | This study |
| NGB467 | *sacA::($P_{veg}$-mCherry, $P_{P1}$-lacIP, $P_{lac}$-yfp) cm, prkC::prkC$^{T314A}$ kan, ΔprpC::tet* | Integration of Δ*prpC::tet* PCR product from JDB2482 genomic DNA using primers NGp67 and NGp196 into NGB458 | This study |
| NGB493 | *sacA::$P_{pdaC}$-luxABCDE cm, prkC::kan* | Transformation of NGB328 into ELB203 | This study |
| NGB494 | *sacA::$P_{pdaC}$-luxABCDE cm, prkC::prkC$^{K40A}$ kan* | Transformation of NGB342 into ELB203 | This study |
| NGB495 | *sacA::$P_{pdaC}$-luxABCDE cm, prkC::prkC$^{T290A}$ kan* | Transformation of NGB345 into ELB203 | This study |
| NGB496 | *sacA::$P_{pdaC}$-luxABCDE cm, prkC::prkC$^{T313A}$ kan* | Transformation of NGB416 into ELB203 | This study |
| NGB497 | *sacA::$P_{pdaC}$-luxABCDE cm, prkC::prkC$^{T320A}$ kan* | Transformation of NGB417 into ELB203 | This study |
| NGB498 | *sacA::$P_{iseA}$-luxABCDE cm, prkC::kan* | Transformation of NGB328 into ELB204 | This study |
| NGB499 | *sacA::$P_{iseA}$-luxABCDE cm, prkC:: prkC$^{K40A}$ kan* | Transformation of NGB342 into ELB204 | This study |
| NGB500 | *sacA::$P_{iseA}$-luxABCDE cm, prkC:: prkC$^{T290A}$ kan* | Transformation of NGB345 into ELB204 | This study |
| NGB524 | *ganA::$P_{xyl}$-prkC$^{T290A\ A292P}$ mls* | Integration of pNG192 into Bs168 | This study |
| NGB527 | *sacA::$P_{pdaC}$-luxABCDE cm, Δ(prpC-prkC)::kan* | Integration of pNG162 into ELB203 | This study |
| NGB528 | *sacA::$P_{pdaC}$-luxABCDE cm, prkC::kan, ΔprpC::tet* | Integration of Δ*prpC::tet* PCR product from JDB2482 genomic DNA using primers NGp67 and NGp196 into NGB493 | This study |
| NGB530 | *Δ(prpC-prkC) sacA::($P_{veg}$-mCherry, $P_{P1}$-lacIP, $P_{lac}$-yfp) cm, ganA::$P_{xyl}$-prkC$^{T290A\ A292P}$ mls* | Transformation of NGB524 into NGB58 | This study |
| NGB531 | *ganA::Pxyl-prkC$^{K40A\ T290A}$ mls* | Integration of pNG193 into Bs168 | This study |

| | | | |
|---|---|---|---|
| NGB535 | Δ(prpC-prkC) sacA::($P_{veg}$-mCherry, $P_{P1}$-lacIP, $P_{lac}$-yfp) cm ganA::$P_{xyl}$-prkC$^{K40A\ T290A}$ mls | Transformation of NGB531 into NGB58 | This study |
| NGB539 | ganA::$P_{xyl}$-prkC-yfp mls | Integration of pNG194 into Bs168. Transcriptional reporter. | This study |
| NGB547 | sacA::$P_{iseA}$-luxABCDE cm, Δ(prpC-prkC)::kan | Integration of pNG162 into ELB204 | This study |
| NGB560 | sacA::($P_{veg}$-mCherry, $P_{P1}$-lacIP, $P_{lac}$-yfp) cm, prkC::prkC$^{K40A}$ kan, ΔprpC::tet, ganA::$P_{xyl}$-prkC$^{K40A\ T290A}$ mls | Integration of pNG207 into NGB382 | This study |
| NGB561 | sacA::($P_{veg}$-mCherry, $P_{P1}$-lacIP, $P_{lac}$-yfp) cm, prkC::kan, ΔprpC::tet, ganA::$P_{xyl}$-prkC$^{K40A\ T290A}$ mls | Integration of pNG207 into NGB379 | This study |
| NGB562 | sacA::($P_{veg}$-mCherry, $P_{P1}$-lacIP, $P_{lac}$-yfp) cm, prkC::prkC$^{T290A}$ kan, ΔprpC::tet, ganA::$P_{xyl}$-prkC$^{K40A\ T290A}$ mls | Integration of pNG207 into NGB380 | This study |

**S3 Table: Plasmids used in this study**

| Plasmid | Genotype | Construction | Source |
|---|---|---|---|
| pNG21 | $P_{veg}$*-mCherry,* $P_{P1}$*-fha2-GGGGSGGGGS-lacI-GSGG-RFTIQEDEEMTKAIPIIKDEE,* $P_{lac}$*-yfp amp cm*, for integration into *sacA* locus | Made by Golden Gate cloning using BsaI. PrkC R280-E300 (RFTIQEDEEMTKAIPIIKDEE) substrate insert amplified from Bs168 genomic DNA with NGp41, NGp42. LacI~P backbone amplified with NGp1, NGp2 from pYW3. Terminator insert amplified with NGp5, NGp6 from pYW1. Referred to throughout as $P_{veg}$*-mCherry,* $P_{P1}$*-lacIP,* $P_{lac}$*-yfp* (see strain table above). | This study |
| pNG130 | $P_{xyl}$*-prkC amp cm*, for integration into *ganA* locus | Made by Golden Gate cloning using BsaI. *prkC* amplified from pEL53 with NGp150, NGp151. Backbone with *Pxyl* promoter amplified from pJMP1 with NGp148, NGp149. | This study |
| pNG131 | $P_{xyl}$*-prkC*$^{T290A}$ *mls amp*, for integration into *ganA* locus | Made by SDM with pNG130 template and mutagenic primers NGp68, NGp69. | This study |
| pNG135 | $P_{xyl}$*-prkC*$^{K40A}$ *mls amp*, for integration into *ganA* locus | Made by SDM with pNG130 template and mutagenic primers NGp152, NGp153. | This study |
| pNG146 | *prkC::kan amp*, for integration into native *prkC* locus | Made by Golden Gate cloning using PaqCI. Vector backbone amplified from pNG130 with NGp173, NGp174. '*rlmN-prpC-prkC-cpgA* from *rlmN(Q30)* amplified from Bs168 genomic DNA with NGp169, NGp170. *kan* cassette amplified from CZ82 with NGp167, NGp168. *thiN'* until *thiN(Q199)* amplified from Bs168 genomic DNA with NGp171, NGp172. | This study |

| | | | |
|---|---|---|---|
| pNG152 | *prkC::prkC*$^{T290A}$ *kan amp*, for integration into native *prkC* locus | Made by SDM with pNG146 template using NGp68, NGp69. | This study |
| pNG154 | *prkC::prkC*$^{K40A}$ *kan amp*, for integration into native *prkC* locus | Made by SDM with pNG146 template using NGp152, NGp153. | This study |
| pNG162 | Δ*(prpC-prkC)::kan amp*, for integration into native *prkC* locus | Made by SDM with pNG146 template using NGp197, NGp198. | This study |
| pNG167 | *P*$_{xyl}$*-prkC*$^{A292P}$ *mls amp* for integration into *ganA* locus | Made by SDM with pNG130 template using NGp200, NGp201. | This study |
| pNG168 | *prkC::prkC*$^{A292P}$ *kan amp*, for integration into native *prkC* locus | Made by SDM with pNG146 template using NGp200, NGp201. | This study |
| pNG171 | *prkC::prkC*$^{I295N}$ *kan amp*, for integration into native *prkC* locus | Made by SDM with pNG146 template using NGp186, NGp187. | This study |
| pNG172 | *prkC::prkC*$^{T313A}$ *kan amp*, for integration into native *prkC* locus | Made by SDM with pNG146 template using NGp202, NGp203. | This study |
| pNG173 | *prkC::prkC*$^{T320A}$ *kan amp*, for integration into native *prkC* locus | Made by SDM with pNG146 template using NGp204, NGp205. | This study |
| pNG177 | *prkC::prkC*$^{T314A}$ *kan amp*, for integration into native *prkC* locus | Made by SDM with pNG146 template using NGp207, NGp208. | This study |
| pNG192 | *P*$_{xyl}$*-prkC*$^{T290A\ A292P}$ *mls amp*, for integration into *ganA* locus | Made by SDM with pNG131 template using NGp201, NGp232. | This study |
| pNG193 | *P*$_{xyl}$*-prkC*$^{K40A\ T290A}$ *mls amp*, for integration into *ganA* locus | Made by SDM with pNG135 template using NGp68, NGp69. | This study |
| pNG194 | *P*$_{xyl}$*-prkC-yfp mls amp,* transcriptional reporter for integration into *ganA* locus | Made by Golden Gate cloning using BsaI. Backbone with *Pxyl* amplified from pNG130 with NGp148, NGp149. *prkC* amplified from pNG130 with NGp150, NGp233. *yfp* insert from pEL53 amplified with NGp234, NGp235. | This study |
| pNG207 | *P*$_{xyl}$*-prkC*$^{K40A\ T290A}$ *mls amp*, for integration into *ganA* locus | Made by transformation of pNG193 into chemically competent 5-alpha. | This study |
| CZ8 | *P*$_{P1}$*-fha2-GSGG-lacI-GSGG-IQEDEEMTKAIPII*, *P*$_{lac}$*-yfp amp cm*, for integration into *sacA* locus | | Lab stock & [9] |
| CZ82 | *P*$_{veg}$*-mCherry kan* | | Lab stock |

| | | | |
|---|---|---|---|
| JN11 | $P_{veg}$-*cfp kan amp* | | Lab stock & [9] |
| pJMP1 | $P_{xyl}$-*dCas9* for integration into *ganA* locus | | Lab stock & Addgene [12] |
| pEB112 | *yfpA206K-FRT-kan-FRT amp* | | Gift from Mark Goulian |
| pDR111 | $P_{hyperspank}$-*MCS spec*, for integration into *amyE* locus | | Lab stock & Addgene [13] |
| pEL53 | $P_{hyperspank}$-*yfpA206K-prkC spec amp*, for integration into *amyE* locus | *prkC* amplified from *B. subtilis* 168 genomic DNA using overlap extension (SOE) PCR with primers nYFP-linker-PrkC and ISp1. Resulting product fused to *yfp* from pEB112 by PCR using primers NheI-RBSYFP-u1 and SphI-prkC-l1 to introduce NheI and SphI restriction sites. Final product digested with SphI and NheI and ligated into pDR111 cut with the same enzymes. | This study |
| pEL186 | $P_{veg}$-*cfp*, $P_{P1}$-*fha2-GSGG-lacI-GSGG-IQEDEEMTKAIPII*, $P_{lac}$-*yfp amp cm*, for integration into *sacA* locus | Made by Golden Gate cloning using BsaI. Backbone amplified from CZ8 with ELp15, ELp16. *Pveg-cfp* insert amplified from JN11 with ELp17, ELp18. | This study |
| pYW1 | $P_{veg}$-*mCherry*, $P_{P1}$-*fha2-GSGG-lacI-GSGG-IQEDEEMTKAIPII*, $P_{lac}$-*yfp amp cm*, for integration into *sacA* locus | Made by Golden Gate cloning using BsaI. Backbone amplified in 2 parts from pEL186 (YWp1, YWp5; YWp2, YWp6), *mCherry* insert amplified from CZ82 with YWp3, YWp4. | This study |
| pYW3 | $P_{veg}$-*mCherry*, $P_{P1}$-*fha2-GGGGSGGGGS-lacI-GSGG-IQEDEEMTKAIPII*, $P_{lac}$-*yfp amp cm*, for integration into *sacA* locus | Made by Golden Gate cloning using BsaI. pYW1 linker extended with mutagenic primers CKp23, CKp24. | This study |

**S4 Table: Oligos used in this study**

| Oligo Name | Sequence (5'-3') |
|---|---|
| NGp1 | GCCGGTCTCAACCCATTAGTTCAACAAACGAAAATTGGATAAAGTGGG |
| NGp2 | TAAGGTCTCAGCCGCCGGATCCCTGCC |
| NGp5 | TTAGGTCTCATGACAGCTTATCATCGGCAA |
| NGp6 | ATAGGTCTCAGGGTGCTTTAGTTGAAGAAT |
| NGp41 | AATGGTCTCACGGCAGATTTACGATTCAAGAAGA |
| NGp42 | GCAGGTCTCAGTCATTCTTCATCTTTAATGATAG |
| NGp64 | TCAAGCCCACGAAGAATG |
| NGp66 | AAACAGGACGCCGTATCAG |
| NGp67 | GGGTGATTGTGGTCGAACTTAG |
| NGp68 | TCTTCTTGAATCGTAAATCTCT |
| NGp69 | TGAAGAAATGGCAAAAGCGATAC |
| NGp100 | CCTTAGGCATGAAAAGCGACGGA |
| NGp148 | ACGGTCTCACTCGAGTAAGGATCTCCAGG |
| NGp149 | CAGGTCTCAGGATCCCATTTCCCCCTTTGATTTTTAGATATCAC |
| NGp150 | CAGGTCTCAATCCATGCTAATCGGCAAGCGGATCAG |
| NGp151 | CAGGTCTCACGAGTTATTCATCTTTCGGATACTCAATGG |
| NGp152 | AGTCGCAATTGCAATCCTGCGG |
| NGp153 | TCACGGTCTAGAATGATATC |
| NGp166 | GTTTTATCCTGAAAATGCG |
| NGp167 | ACCACCTGCAATACTGTAGAAAAGAGGAAGG |
| NGp168 | AGCACCTGCGGTACTAAAACAATTCATCCAG |
| NGp169 | AACACCTGCTGGGTGGGCAATGGCTGACAGACAATGG |
| NGp170 | GACACCTGCTGGGACAGTTATTTACTTCCTCTGATTTCAGAAATTGC |
| NGp171 | AGCACCTGCTTAGTTAGGCAGCTTTAGAAAAGAGCTGGG |
| NGp172 | AACACCTGCTGGGTGGGCCTCGTGAATGAATGAGTTCG |
| NGp173 | TACACCTGCTCACCCCATCCTGATATTGTCTGC |
| NGp174 | GACACCTGCTTAGCCCATCATTCTTGAAGACG |
| NGp175 | CAATGACAAGACCGATACCTATCATTAAAGATGAAG |
| NGp184 | GGCACTCTTGTCTACAGTACG |
| NGp185 | ATCGCCCGGACAATTCCTCC |
| NGp186 | GCGATACCTAACATTAAAGATG |
| NGp187 | TTTTGTCATTTCTTCATCTTC |
| NGp196 | ATCGGTGCTCGCCATATTACGG |
| NGp197 | CAGCATCTCTGACACTTACG |
| NGp198 | AATCGTCTCTGCTTCATTCC |
| NGp200 | AATGACAAAACCGATACCTATCATTAAAG |
| NGp201 | TCTTCATCTTCTTGAATCG |
| NGp202 | AGCTGAAGTGGCAACCGCACA |
| NGp203 | TCTTTTTCGCCAGCAGCTTTAG |
| NGp204 | AGAAAACAAAGCAAAGAAGAACGG |
| NGp205 | TGTGCGGTTGTCACTTC |
| NGp207 | TGAAGTGACAGCAGCACAAGAAAAC |
| NGp208 | GCTTCTTTTTCGCCAG |
| NGp232 | AATGGCAAAACCGATACCTATC |
| NGp233 | CGGTCTCGATCTCCTTCTTATTCATCTTTCGGATACTCAATGG |

| | |
|---|---|
| NGp234 | AAGGTCTCGAGATACCATGCGTAAAGGAGAAGAACTTTTCACTGG |
| NGp235 | AGGTCTCTCGAGTTATTTGTATAGTTCATCCATGCCATGTGTAATCCC |
| CKp21 | ATCATTGTTTCCCCCGCCAT |
| CKp22 | GTCATGGGGGCCAGCTATAC |
| CKp23 | GGTCTCAAAGCGGAGGTGGAGGAAGCAAACCAGTAACGTTATACGATG |
| CKp24 | GGTCTCAGCTTCCACCACCGCCTCCAAGCTTTTTAACGAGGTC |
| ELp15 | GGCTACGGTCTCGCTTAAGCATCACCGGCGCCACAG |
| ELp16 | GGCTACGGTCTCGACGATGCGTCCGGCGTAG |
| ELp17 | GGCTACGGTCTCGTCGTGGCCGAATTTTGTCAAAATAATTTTATTGACAAC |
| ELp18 | GGCTACGGTCTCGTAAGTGCCCTTATACAACTC |
| nYFP-linker-PrkC | GGCATGGATGAACTATACAAATCAGGAGGCTCTCTAATCGGCAAGCGGATCAGC |
| ISp1 | GGGCATGCGCCGCTTAGCGCCTT |
| NheI-RBSYFP-u1 | GGCTAGCTAGCAAAAGGTGGTGAACTACTATGCGTAAAGGAGAAGAACTTTTC |
| SphI-prkC-l1 | GGCTAGCATGCTTATTCATCTTTCGGATAC |
| KVp18 | CCGAAGGTGAGCCAGTGTGACT |
| YWp1 | GAGGTCTCGCATCACCGGCGCCACAGGT |
| YWp2 | AGGGTCTCTTAATCGCTAGCTTTTCTCCACATTTATTGTACAACACGAG |
| YWp3 | GCGGTCTCGATTAACTAATAAGGAGGACAAACATGGTTTCCAAGGGCGAGGAGG |
| YWp4 | GAGGTCTCTGATGCTTATTTGTACAGCTCATCCATGCCACC |
| YWp5 | ATGGTCTCTCCCGCCGTTAACCACCATCAAACAG |
| YWp6 | ATTGGTCTCGCGGGATATAACATGAGCTGTCTTCG |
| YWp18 | CTCCTTTGTTTATCCACCGAAC |

#### Supplementary References


1 Stancik, I. A. *et al.* Serine/Threonine Protein Kinases from Bacteria, Archaea and Eukarya Share a Common Evolutionary Origin Deeply Rooted in the Tree of Life. *J Mol Biol* **430**, 27-32 (2018). https://doi.org/10.1016/j.jmb.2017.11.004

2 Jumper, J. *et al.* Highly accurate protein structure prediction with AlphaFold. *Nature* **596**, 583-589 (2021). https://doi.org/10.1038/s41586-021-03819-2

3 Abramson, J. *et al.* Accurate structure prediction of biomolecular interactions with AlphaFold 3. *Nature* **630**, 493-500 (2024). https://doi.org/10.1038/s41586-024-07487-w

4 Consortium, T. U. UniProt: the Universal Protein Knowledgebase in 2025. *Nucleic Acids Research* **53**, D609-D617 (2024). https://doi.org/10.1093/nar/gkae1010

5 Hallgren, J. *et al.* DeepTMHMM predicts alpha and beta transmembrane proteins using deep neural networks. *bioRxiv*, 2022.2004.2008.487609 (2022). https://doi.org/10.1101/2022.04.08.487609

6 Altschul, S. F. *et al.* Gapped BLAST and PSI-BLAST: a new generation of protein database search programs. *Nucleic Acids Res* **25**, 3389-3402 (1997). https://doi.org/10.1093/nar/25.17.3389

7 Jones, P. *et al.* InterProScan 5: genome-scale protein function classification. *Bioinformatics* **30**, 1236-1240 (2014). https://doi.org/10.1093/bioinformatics/btu031

8 Harwood, C. R. & Cutting, S. M. *Molecular biological methods for Bacillus*. Vol. 647 (Wiley New York, 1990).

9 Zheng, C. R., Singh, A., Libby, A., Silver, P. A. & Libby, E. A. Modular and Single-Cell Sensors of Bacterial Ser/Thr Kinase Activity. *ACS Synthetic Biology* **10**, 2340-2350 (2021). https://doi.org/10.1021/acssynbio.1c00250

10 Libby, E. A., Reuveni, S. & Dworkin, J. Multisite phosphorylation drives phenotypic variation in (p)ppGpp synthetase-dependent antibiotic tolerance. *Nature Communications* **10**, 5133 (2019). https://doi.org/10.1038/s41467-019-13127-z

11 Libby, E. A., Goss, L. A. & Dworkin, J. The Eukaryotic-Like Ser/Thr Kinase PrkC Regulates the Essential WalRK Two-Component System in Bacillus subtilis. *PLOS Genetics* **11**, e1005275 (2015). https://doi.org/10.1371/journal.pgen.1005275

12 Peters, J. M. *et al.* A Comprehensive, CRISPR-based Functional Analysis of Essential Genes in Bacteria. *Cell* **165**, 1493-1506 (2016). https://doi.org/https://doi.org/10.1016/j.cell.2016.05.003

13 Ben-Yehuda, S., Rudner, D. Z. & Losick, R. RacA, a bacterial protein that anchors chromosomes to the cell poles. *Science* **299**, 532-536 (2003). https://doi.org/10.1126/science.1079914
